# Feasibility of methionine oxidation quantification in proteins through Raman and FT-IR spectroscopy

**DOI:** 10.64898/2026.11.18.767667

**Authors:** Thorben Höltkemeier, Ingo Fischer, Marco Venerito, Victor Le-Minh, François Griaud, Patrick Merkle, Manuela Gritsch, Adam Sobczuk, Malak Bitar, Eva Krois, Patrick Kubryk, Karoline Bechtold-Peters, Christoph Haisch, Wolfgang Friess

## Abstract

The development of biopharmaceuticals poses unique challenges, with alterations of chemical integrity potentially impairing product efficacy and safety. Thus, oxidation-susceptibility of drug candidates is typically assessed by chromatographic methods, potentially coupled with mass spectrometry (LC-MS), which enable residue-specific oxidation monitoring but require high-resolution instrumentation, specialized expertise, and may introduce sample preparation artefacts. These constraints have raised interest in orthogonal, simple and fast methods to probe oxidation in native protein samples. Raman and Fourier transform infrared spectroscopy (FT-IR) have shown potential for detecting methionine oxidation, the most common oxidative protein liability. The aim of this work was to transfer the known quantification potential of cuvette-based Raman spectroscopy to Raman microscopy and evaluate its suitability for high-throughput screenings. Following several technical optimizations, this work shows that the detection of oxidation by Raman microscopy could only be confirmed for molecular methionine in solution and not for proteinogenic methionine. In contrast, liquid-cell FT-IR enabled solid quantitative read-outs of methionine oxidation in proteins. Under the investigated forced degradation conditions, the method showed good agreement to orthogonal LC-MS measurements utilizing the multi-attribute method (MAM). These findings position liquid-cell FT-IR as a potential rapid, low-volume screening tool for oxidation assessments in native protein samples, particularly during early-stage biopharmaceutical development.

## 1 Introduction

Protein pharmaceuticals have become an established therapeutic platform. Offering advanced selectivity, well-studied safety profiles as well as rational design opportunities, they are favorable drug candidates in a number of areas ranging from cancer and autoimmune to degenerative diseases. While monoclonal antibodies (mAb) are a major platform, novel protein formats are emerging(de la Torre and Albericio, 2022; Kesik-Brodacka, 2018). This increased diversity comes with the need for a deeper understanding of potential molecule instabilities, referred to as liabilities, already in early development (Condado-Morales et al., 2024). Among those vulnerabilities, chemical integrity plays a key role in maintaining functionality and structure. The most prevalent chemical changes observed in biopharmaceutical development are deamidation, isomerization and oxidation. Deamidation and isomerization are solvent-catalyzed, pH-dependent reactions that can be addressed by the selection of a suitable pH (Lu et al., 2019; Torosantucci et al., 2014; Zeunik et al., 2022). In marketed biopharmaceutical products, oxidation most commonly affects methionine (Met) residues, forming methionine-sulfoxide (MetOx), followed by the oxidation of tryptophan, cysteine, histidine, and tyrosine(Le Basle et al., 2020; Torosantucci et al., 2014). Contaminants such as peroxides, metals or molecular oxygen mediate the oxidation reaction. Reactive oxygen species can originate from contaminations in excipients such as polysorbates, process-related sources like sterilization protocols, including vaporized hydrogen peroxide, form upon light exposure or from metal contamination (Meleties et al., 2023; Sreedhara et al., 2016, 2013; Subelzu and Schöneich, 2021; Wuchner et al., 2022). Because contaminant concentrations are usually low, effects often only show after longer storage times.

In drug development, oxidation liability is traditionally assessed by forced degradation studies (FDS), exposing the drug candidate to excess amounts of an oxidant such as hydrogen peroxide or 2,2’-azobis (2-amidinopropane) dihydrochloride (AAPH) (Halley et al., 2020; Heinzl et al., 2019). Oxidative changes can be examined by a range of methods. Standard FDS protocols typically include analysis of oxidative changes based on orthogonal analytical methods such as capillary electrophoresis, ion-exchange chromatography and LC-MS as well as methods that assess secondary effects of oxidation, such as reversed phase HPLC (RP-HPLC), size exclusion chromatography (SEC), or particle counting (Halley et al., 2020; Hipper et al., 2022). Peptide mapping methods enable sensitive and site-specific detection of peptide oxidation using LC-MS instrumentation after enzymatic digestion. The multi-attribute method (MAM) applies a peptide library for consistent monitoring of post-translational modifications characterized by tandem LC-MS/MS data acquired from degraded samples. Subsequent LC-MS acquisitions and comparison with the established library allow for a consistent detection of multiple post-translational modifications, including oxidation on the level of single amino acid residues (site-specific detection) in any protein format. The method is highly sensitive and typically provides relative estimates of oxidation levels (Pohl et al., 2023; Rogers et al., 2015; Yang et al., 2023). MAM requires access to complex and costly LC-MS instrumentation and MS expertise to enable library generation and data interpretation. Moreover, a robust automation platform is required to control potential artefacts introduced during sample preparation. (Griaud et al., 2026; Mouchahoir and Schiel, 2018). To reduce the experimental workload in oxidation assessment, various *in silico*-based approaches for predicting oxidation liabilities have been introduced in recent years. This includes machine learning models building on experimental data derived from FDS as well as structure-based predictions focusing on the correlation of solvent accessible surface area (SASA) of individual residues with their susceptibility to oxidation (Delmar et al., 2021; Sankar et al., 2018; Tavella et al., 2022; Yang et al., 2017). Homology-based models are usually focused on antibodies and are not transferable to novel protein formats, while universally applicable models are limited to the prediction of qualitative likelihood of oxidation and do not enhance the understanding of the quantitative extent of oxidation under specific conditions. The need for experimental verification and quantification of predicted oxidation events under FDS or process-relevant conditions demands access to simple and fast alternatives to MAM. Alternatives focus on changes in chromatographic or electrophoretic behavior of an oxidized molecule, including affinity RP-HPLC and capillary zone electrophoresis (Loew et al., 2012; Mo et al., 2016; Saunders and Stites, 2012; Schöneich, 2005). These methods probe changes in the physico-chemical behavior of molecules and can only distinguish whether observed species are related to Met oxidation or other modifications when accompanied by orthogonal characterization. In comparison, vibrational spectroscopy directly assesses changes in chemical bonds. Hence, Raman and Fourier transform infrared spectroscopy (FT-IR) have been proposed as valid alternatives for specific detection of Met oxidation (Balakrishnan et al., 2018; Ravi et al., 2011; Torreggiani et al., 2011). Balakrishnan et al. (2018) showed the applicability of cuvette-based Raman spectroscopy for quantification of Met oxidation in highly concentrated protein samples. FT-IR in ATR mode has been utilized for qualitative detection of oxidative changes in proteinogenic and molecular Met (Balakrishnan et al., 2018; Ravi et al., 2011; Torreggiani et al., 2011).

This work aimed to transfer the usage of Raman spectroscopy for Met oxidation detection to Raman microscopy to enable automated, small-volume, high-throughput screenings that are not feasible with cuvette-based systems. These capabilities sought to be enabled using sample preparation in closed micro chambers and relying on the microscopic system for automated acquisition. For this purpose, the optimal technical set-up in terms of laser wavelength and power, grating and objective for measurement of protein solutions was identified. These optimizations enabled the detection, linearity assessment and quantification of MetOx in solution. Nonetheless, this work showed that results from cuvette-based Raman spectroscopy were not transferable to Raman microscopy set-ups, as the relevant spectral changes were not observed in oxidized proteinogenic Met. In contrast, liquid-cell FT-IR did enable quantification of the bulk degree of Met oxidation with results showing agreement to orthogonal MAM measurements. This work builds on the specific S=O vibrational band at 1009 cm^-1^ that appears upon formation of MetOx and showed linear correlation with molecular and proteinogenic MetOx. The evaluation of method performance focused on method development and feasibility rather than formal analytical procedure validation. Based on the calibration, liquid-cell FT-IR quantified Met oxidation with low measurement variability in two model proteins subjected to different oxidative stresses. Therefore, liquid-cell FT-IR presented its potential as a rapid, low-volume, non-invasive screening tool for mid-to high concentration protein formulations in early phases of drug development.

## 2 Materials and Methods

### 2.1 Materials

DL-methionine, L-methionine-sulfoxide, and 30% Hydrogen peroxide were obtained from Sigma-Aldrich (St. Louis, Missouri, USA). Human Serum Albumin (HSA; Sigma-Aldrich) and lysozyme (from chicken egg white, Fluka Analytical, Sigma-Aldrich) were utilized as reference molecules. Additional excipients included sucrose, sodium chloride, and polysorbate 20 and 80 of at least a “Ph.Eur.” grade. All solutions were prepared in Milli-Q water (Millipore Integral 3^TM^, Merck). Chemical denaturation was achieved by dilution in 8 M guanidinium hydrochloride solution at pH 8.5 (Sigma-Aldrich). Pharmaceutical drug products included monoclonal antibodies (mAbs) and non-antibody therapeutic proteins (TPs) at concentrations of up to 150 mg/ml formulated in histidine, citrate, or Tris buffer.

Mass spectrometry grade deionized water was provided by a Milli-Q system (Merck Millipore; Burlington, MA, USA). Acetonitrile (LC-MS grade) and iodoacetamide were obtained from Merck KGaA (Darmstadt, Germany). Pierce dithiothreitol and trifluoroacetic acid were obtained from Thermo Fisher Scientific (Waltham, MA, USA). Lysyl endopeptidase (LysC, MS grade) was obtained from Fujifilm Wako Pure Chemical (Osaka, Japan), while trypsin gold (MS grade) was purchased from Promega (Madison, WI, USA).

### 2.2 Sample preparation

Dilution series of Met and MetOx were prepared from a 100 mM stock solution in water. Physical mixtures of Met/MetOx with proteins were prepared in matching formulations. Proteins were exposed to oxidative stress through the addition of hydrogen peroxide at a final concentration of 0.1 to 1000 mM and incubation at 25 °C in the dark for 24 hours. Cut pieces of catalysator-coated plastic rings from disinfection sets for contact lenses (e.g. AOSept Plus Hydraglyde, Alcon Laboratories, Inc, Fort Worth, TX, USA) were used as catalysts for quenching by incubation at room temperature for 2 hours. In studies with free Met as antioxidant, samples were buffer exchanged in Amicon spin columns (Amicon ^®^Ultra 15 mL Centrifugal Filters Ultracel ^®^ -30K; Sigma-Aldrich) to remove free MetOx.

### 2.3 Microscope-based Raman Spectroscopy

Raman spectra were acquired using three Raman microscopes. System A was a WiTec ALPHA300 R Confocal Raman microscope system (Abingdon, UK; Oxford Instruments formerly known as WITec GmbH, Ulm, Germany) equipped with an excitation wavelength of 532 nm Nd:YAG laser (WITec, Ulm, Germany) and Zeiss LD Plan-Neofluar 63× / 0.75, Zeiss LD Plan-Neofluar 40× / 0.6, Zeiss EC Epiplan-Neofluar HD 20× / 0.5, Zeiss Plan-Apochromat SF25 20× / 0.8, Zeiss EC Epiplan-Neofluar Pol 5× / 0.13 microscope objectives. Raman scattering was collected using a UHTS 300 VIS spectrometer and a charge-coupled device (CCD) camera with G1: 1800 g/mm BLZ 500nm grating applied, and a spectral center of 700.0 cm^-1^. An offset correction was performed for each study by calibration against a silicon substrate (520.7 cm^-1^). The excitation power before the objective was 68 mW, the integration time was 600.0 s, and a single scan recorded unless indicated otherwise. Cosmic ray removal and baseline correction of Raman spectra were performed using WiTec Project Plus 5.3 software.

System B was a WiTec ALPHA300 R system from Oxford instruments (Abingdon, UK; formerly known as WITec GmbH, Ulm, Germany). It was equipped with a Cobolt DPL 532 nm solid-state laser (Cobolt AG, Solna, Sweden) and a True Power module. A 50× / 0.55 objective from Carl Zeiss AG was used for all measurements. The system contained a spectrometer with a grating of 600 g/mm and a Newton 970 electro-multiplied charge-coupled device (EMCCD) camera (Andor Technology Ltd, Belfast, UK). Spectra of two-second integration time and up to 30 accumulations were recorded with 50 mW laser power.

For measurements with system B, 140 µm thick double-sided adhesive tape (ARcare® 90106NB from Adhesive Research Inc., Glen Rock, PA, USA) was cut into 25.0 mm × 25.0 mm squares, containing a circular cavity with a diameter of 13.0 mm in the middle of it. This tape was placed onto a microscope slide (Thermo Scientific, Waltham, MA, USA), which was cleaned with water and ethanol prior to use. 10 µL of the sample was placed into the cavity, and the system was closed with a pre-cleaned (rinsed with water and ethanol) 18.0 mm × 18.0 mm coverslide (Marienfeld Superior, Lauda-Königshofen, Germany). Excess liquid was removed with a tissue, and the closed chamber was sealed with wax to prevent evaporation of water during the measurements.

System C was an inVia Qontor from Renishaw plc (Wotton-under-Edge, UK) equipped with 457 nm, 532 nm, 633 nm, and 785 nm lasers (50 mW, 100 mW, 16 mW, and 300 mW, respectively) and a Leica N PLAN 20× / 0.40 microscope objective. A CCD camera and gratings with 600 g/mm, 1200 g/mm, 1800 g/mm or 2400 g/mm were used to collect the Raman signal. An integration time of 200.0 s was used. The available technical components differed for the three systems and are summarized in Table 1.

**Table 1.**
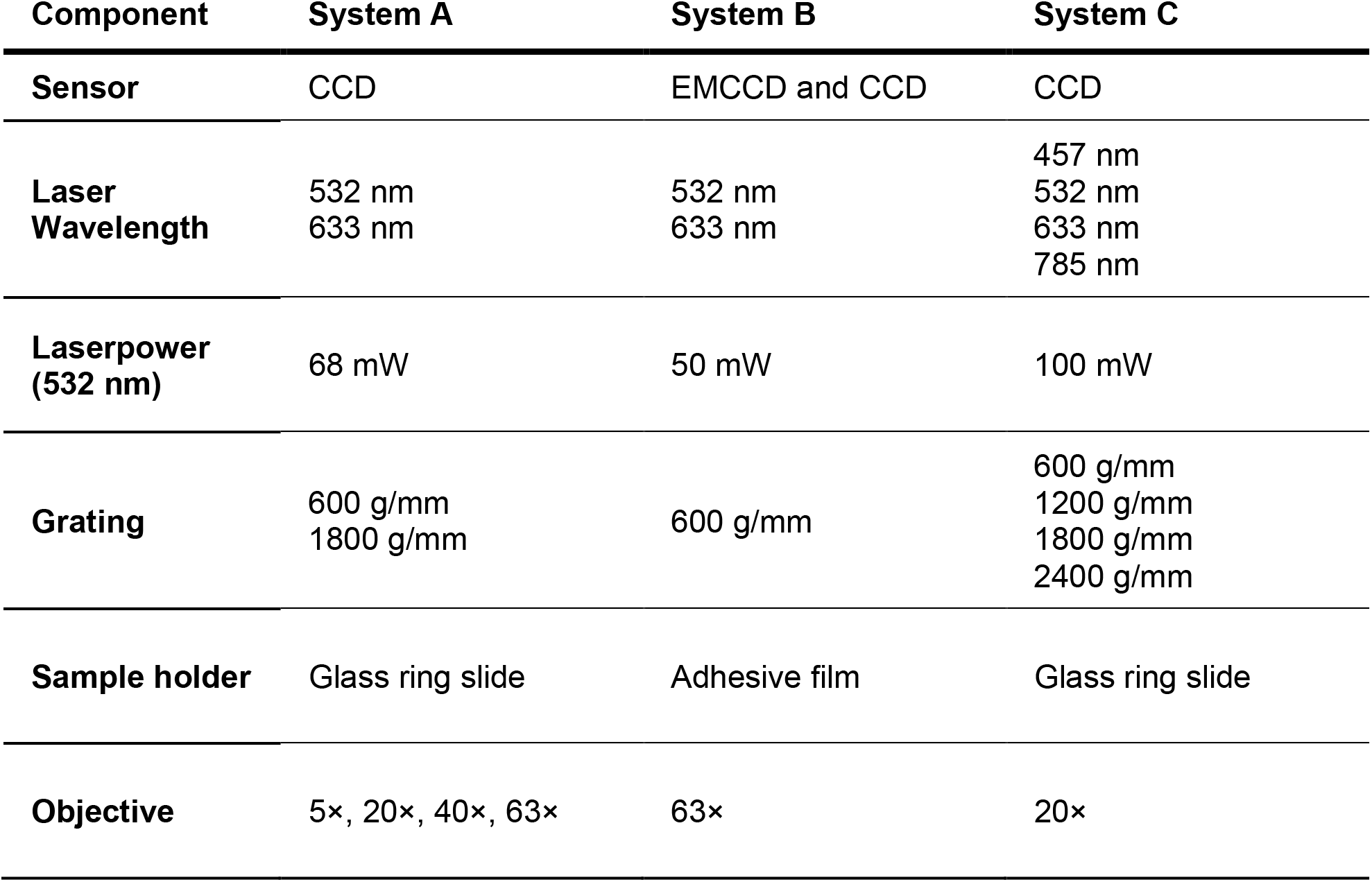
Technical components available in different Raman systems used for this study.

| Component | System A | System B | System C |
|---|---|---|---|
| **Sensor** | CCD | EMCCD and CCD | CCD |
| **Laser Wavelength** | 532 nm<br>633 nm | 532 nm<br>633 nm | 457 nm<br>532 nm<br>633 nm<br>785 nm |
| **Laserpower (532 nm)** | 68 mW | 50 mW | 100 mW |
| **Grating** | 600 g/mm<br>1800 g/mm | 600 g/mm | 600 g/mm<br>1200 g/mm<br>1800 g/mm<br>2400 g/mm |
| **Sample holder** | Glass ring slide | Adhesive film | Glass ring slide |
| **Objective** | 5×, 20×, 40×, 63× | 63× | 20× |

### 2.4 Fourier-transform Infrared Spectroscopy (FT-IR)

FT-IR spectra were recorded in a Bruker Tensor II system (Bruker, Billerica, MA, USA) equipped with an AquaSpec cell, flow-through QL (Type: A741/Q) in the wavenumber range of 950 to 3100 cm^-1^ with a 0.71 cm^-1^ interval. Spectra were averaged from 60 scans (background) or 120 scans (sample). A 50 µl Hamilton glass syringe (Type 1705N) was washed with the sample solution for equilibration. The measurement cell was flushed through with the injection of 20 µl sample volume. Three replicates were measured per sample by injection of 10 µl each. Syringe and cell were cleaned in between each sample with three times 50 µl Milli-Q water.

### 2.5 LC-MS -Multi Attribute Method

Protein samples were digested into peptides using a STARLET robotic system (Hamilton, Bonaduz, Switzerland) equipped with a TRobot II PCR thermal cycler (Biometra, Analytik Jena, Jena, Germany) as previously described (Griaud et al., 2026). First, disulfide bonds were reduced using 5 mM DTT in 4.7 M Guanidine hydrochloride (Sigma-Aldrich), followed by cysteine carbamidomethylation with 9 mM IAM. Enzymatic proteolysis was then carried out using a combination of LysC and trypsin at 37 °C for 20 hours. Protein digestion was subsequently quenched by acidifying with 0.1% (v/v) trifluoroacetic acid (TFA) prior to LC-MS analysis.

Resulting peptide mixtures (4 µg) were separated on an Acquity HSS T3 column (100 Å, 1.7 µm, 2.1 mm x 150 mm). In a mobile phase containing 0.1 % TFA the 69 min linear gradient moved from 10 to 47.5 % of the secondary mobile phase containing 0.09 % TFA in acetonitrile. The protocol was established on a Thermo Vanquish UHPLC system coupled to a Thermo Orbitrap Exactive Plus mass spectrometer (Thermo Fischer Scientific, MA, US).

The mass spectrometer was equipped with a heated electrospray ionization source. In positive ionization mode, a spray voltage of 3.7 kV and a temperature of 250 °C were selected. Sheath gas flow and aux gas flow were set at 45 and 10 (arbitrary units), respectively. Capillary temperature was set to 250 °C. Mass spectrometry data were processed using Genedata Expressionist software v17.5 to perform targeted identification of native peptides and related oxidized species based on their mass-to-charge ratio and retention time as previously described (Griaud et al., 2026).

### 2.6 Methionine abundance and oxidation liability

The total abundance of Met residues in a molecule was determined from the sequence. Residues exhibiting a relative SASA ≥ 15 % were labeled as liable (Tavella et al., 2022). SASA was determined by PyMOL (Version 2.5.0 Schrödinger, LLC.). Structures were based on internal crystal structures, homology modeling or AlphaFold2; crystal structures of HSA, lysozyme and NIST-Fab were obtained from RCSB.org (PDB IDs: 1AO6, 168L, 5K8A).

## 3 Results

The potential quantification capabilities of both Raman and FT-IR were evaluated in three steps. First, the spectral changes caused by oxidation of free Met in solution were assessed for qualitative identification. Secondly, the concentration-dependent linearity of the observed spectral changes was evaluated based on free MetOx to calibrate the system for quantification. Finally, the agreement of the method with orthogonal, semi-quantitative MAM results was assessed.

### 3.1 Methionine oxidation detection in Raman spectroscopy

Oxidation of free Met altered four bands of its Raman spectrum (Figure 1 a). Two distinct bands appeared at 945 and 1009 cm^-1^, which are related to the S=O bond of sulfoxide. The CH_2_ band at 1428 cm^-1^ and the CS band at 702 cm^-1^ increased in intensity (Figure 1 a). In proteins, these bands were all overlaid by the background spectrum of the molecule except for the CS band at 702 cm^-1^. With Raman system A, the peak height of MetOx at 702 cm^-1^ and Met at 699 cm^-1^ and 722 cm^-1^ scaled linearly with concentration (Figure 1 b). The LOD was evaluated based on signal-to-noise ratio (SNR) and on the standard deviation of a linear response and a slope following ICH Q2(R2) guidelines (ICH, 2023). Both approaches build on the standard deviation of the background *s_B_*, that was calculated based on the water baseline (n = 10) at the respective Raman shift. The SNR approach (further referred to as 3-*s_B_* criterion) defines the LOD at a SNR of 3:1 which corresponded to 3.0 ± 0.3 mM MetOx in system A and 2.8 ± 0.4 mM MetOx in system C. Met calibration in system A at 699 cm^-1^ exhibited an LOD of 6.6 ± 1.1 mM. The second approach calculates LOD from the slope of the calibration curve *S* and the standard deviation of the response *σ*. The value of *σ* can be estimated based on the standard deviation of the background (*σ* = *s_B_*) leading to Equation 1.

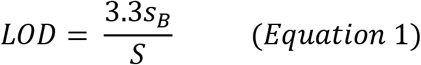

**Figure 1.**
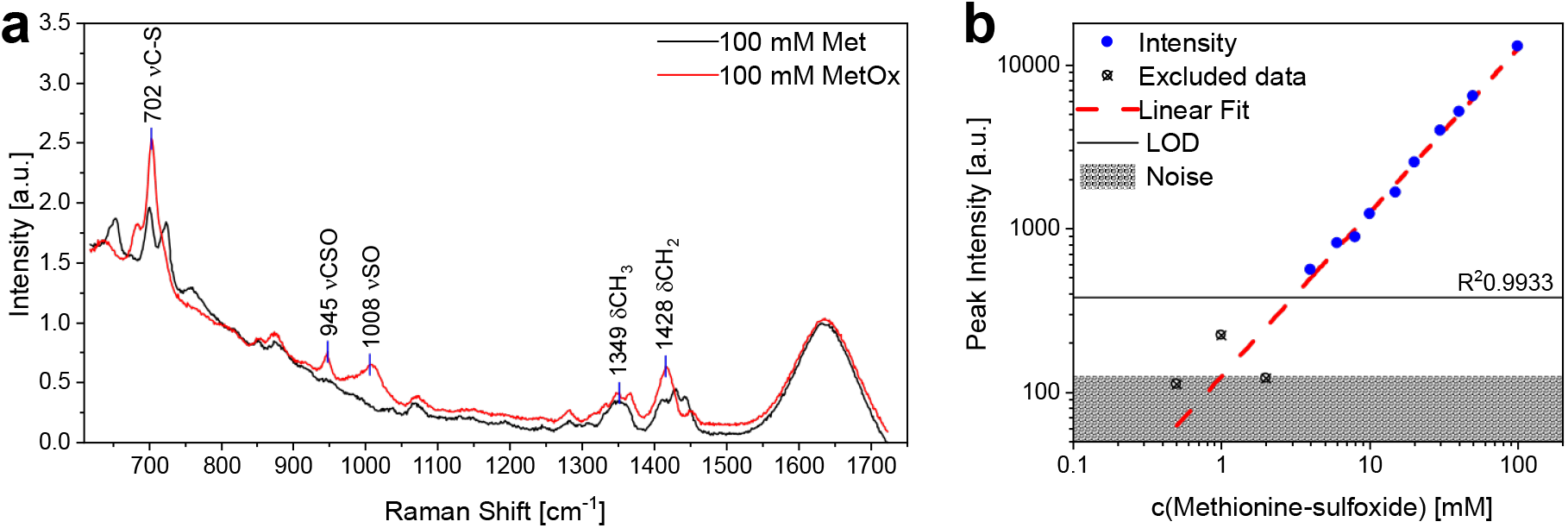
Detection of Met oxidation in solution using Raman spectroscopy; (a) Raman spectra of Met and MetOx with peak annotations (n = 1; system A, 532 nm laser, 20x objective, 600 s integration time) ; (b) calibration of system A regarding peak intensity at 702 cm^-1^ of molecular MetOx in aqueous solution, Noise represents the standard deviation of n = 10 water baseline measurements at 702 cm^-1^, LOD is estimated based on the 3-s_B_ criterion and found to be 3.0 ± 0.3 mM MetOx.

This LOD approximation resulted in values of 3.3 ± 0.03 mM MetOx in system A, 1.8 ± 0.06 mM MetOx in system C and 8.8 ± 0.2 mM Met in System A. Raman system B was not evaluated for linearity of detection or LOD.

### 3.2 Method optimization for methionine oxidation detection in Raman spectroscopy

To achieve maximum SNR for Met oxidation analysis in Raman spectroscopy, eight potential influencing factors were studied. In presence of analyte, noise can be estimated from regions of the baseline adjacent to the peak of interest (ICH, 2023). Thus, it was approximated based on the standard deviation *s_Noise_* of all data points (n = 69) in the range between 640 and 600 cm^-1^, as well as 740 and 800 cm^-1^, after subtraction of the matching buffer reference. The signal intensity was measured at 702 cm^-1^ after reference subtraction and set in reference to 3 ∗ *s_Noise_*.

Neither sensor type (CCD or EMCCD), sample holder (ring slide or adhesive film), nor the usage of stainless-steel mirror finish or gold-coated slides to trigger surface-enhanced Raman scattering (SERS) improved the SNR. In set-up A, with different objectives, a 20x 0.5 dark field (DF) showed the highest sensitivity. The choice of grating is laser dependent but due to higher resolution and better SNR, the highest grating number feasible at the chosen wavelength is desired. An increase in laser power improved SNR. No loss of signal was observed over prolonged measurements of up to 12 h in protein or free Met samples, indicating that the samples stayed stable under measurement conditions and continuous laser exposure. The 532 nm laser exhibited the best SNR (Figure 2 a) in set-up C while also having the highest power output (100 mW). The most significant increase in SNR was achieved by an increase of measurement time. Comparable levels of SNR were reached at any total measurement time independent of the combination of accumulations and integration time used (Figure 2 b). The increase of SNR was observed to be proportional to the measurement time squared (*SNR* ∝ *t_measurement_*^2^) in the observed range. Thus, no substantial gain in SNR is expected beyond a total measurement time of ∼ 600 s.

**Figure 2.**
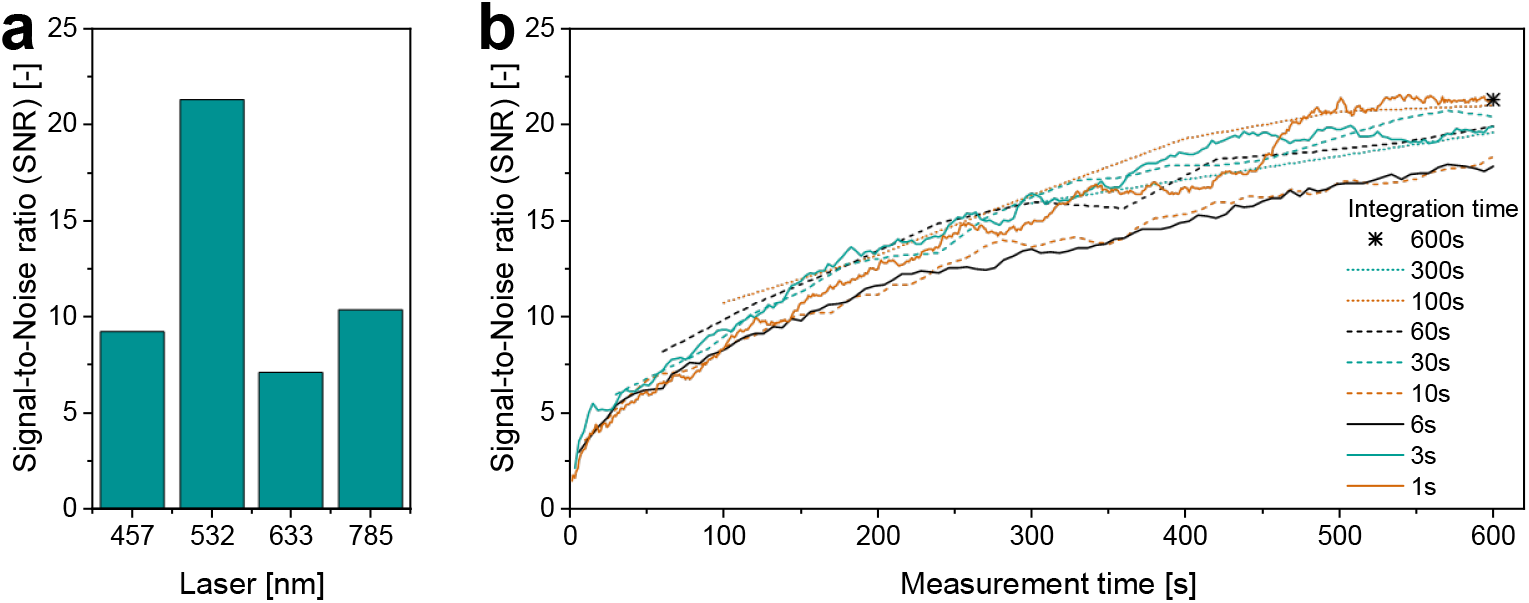
Signal-to-noise ratio (SNR) of the C-S band at 702 cm^-1^ in Raman spectroscopy; (a) SNR of four different laser wavelengths in Raman system C based on 250 mg/ml lysozyme solution; (b) SNR of 50 mM MetOx in aqueous solution in relation to total measurement time in Raman system A (532 nm laser at 68 mW, 1800 g/mm, 20x 0.5 objective), at any total measurement time all available accumulations with the set integration time were averaged, all accumulations at a set integration time were measured in a single run.

### 3.3 Detection and quantification of methionine oxidation in proteins in Raman spectroscopy

Three mAbs and two TPs at 10 to 90 mg/ml were exposed to up to 1000 mM H_2_O_2_ and screened for changes of their Raman spectra at 702 cm^-1^. None of the samples yielded a relevant read-out in this region within a measurement time of up to 2 h (Figure 3, exemplary for mAb1; other proteins see supplementary data). An increase in concentration of mAb1 up to 150 mg/ml and a measurement time of up to 2 h did not result in significant changes at 702 cm^-1^ (Figure 3 b). In an effort to increase detection accessibility, the chemical denaturation with 4.5 and 6 M guanidinium hydrochloride did not lead to the detection of Met oxidation in mAb2 at 54 mg/ml after exposure to 1000 mM H_2_O_2_.

**Figure 3.**
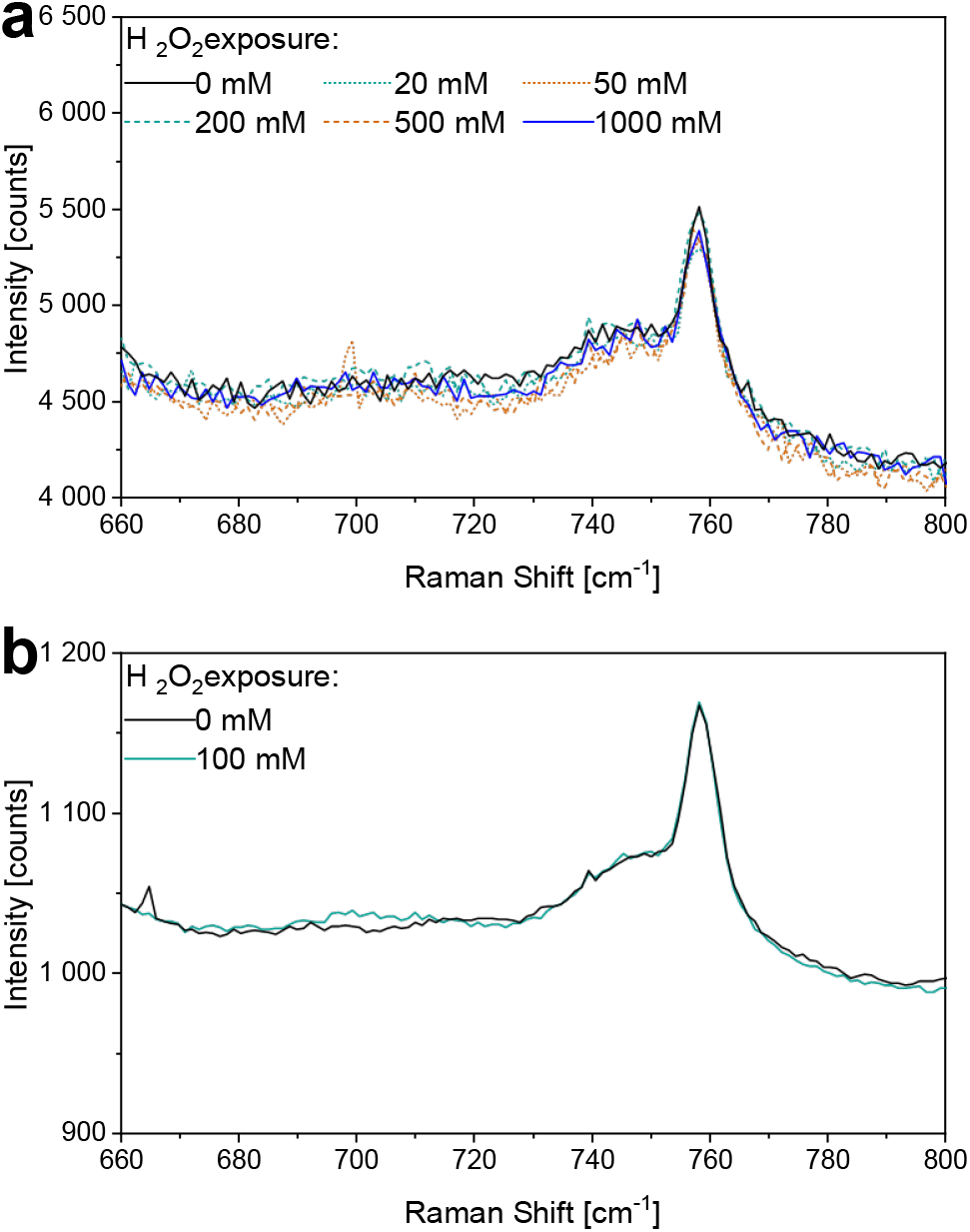
Baseline corrected Raman spectra of mAb1 under varying oxidative stress in system A; (a) protein concentration 90 mg/ml, total measurement time 10 min (1 x 600 s), settings: CCD detector, 1800 g/mm grating, 532 nm laser at 68 mW, 20x objective; (b) protein concentration 150 mg/ml, total measurement time 2h (120 x 60 s), settings: CCD detector, 1800 g/mm grating, 532 nm laser at 68 mW, 20x objective.

### 3.4 Methionine oxidation in FT-IR spectroscopy – Peak identification

Oxidation of molecular Met affected four bands in the FT-IR spectrum in the range of 950 to 2000 cm^-1^. The bands at approx. 1345 and 1443 cm^-1^ decreased in intensity while two new bands at around 1330 and 1009 cm^-1^ appeared (Figure 4 a). The relevant marker for oxidation detection is the S=O vibrational band at 1009 cm^-1^. This band overlaps with glycosylation related bands in therapeutic proteins as demonstrated for mAb3, which has a standard glycosylation pattern and HSA which is not glycosylated (Figure 4 b + c). The overlap with glycosylation caused a slight shift of the peak maximum down to 1004 cm^-1^, which was absent after subtraction of the protein signal. The overlap of protein and MetOx signal was additive in mixtures.

**Figure 4.**
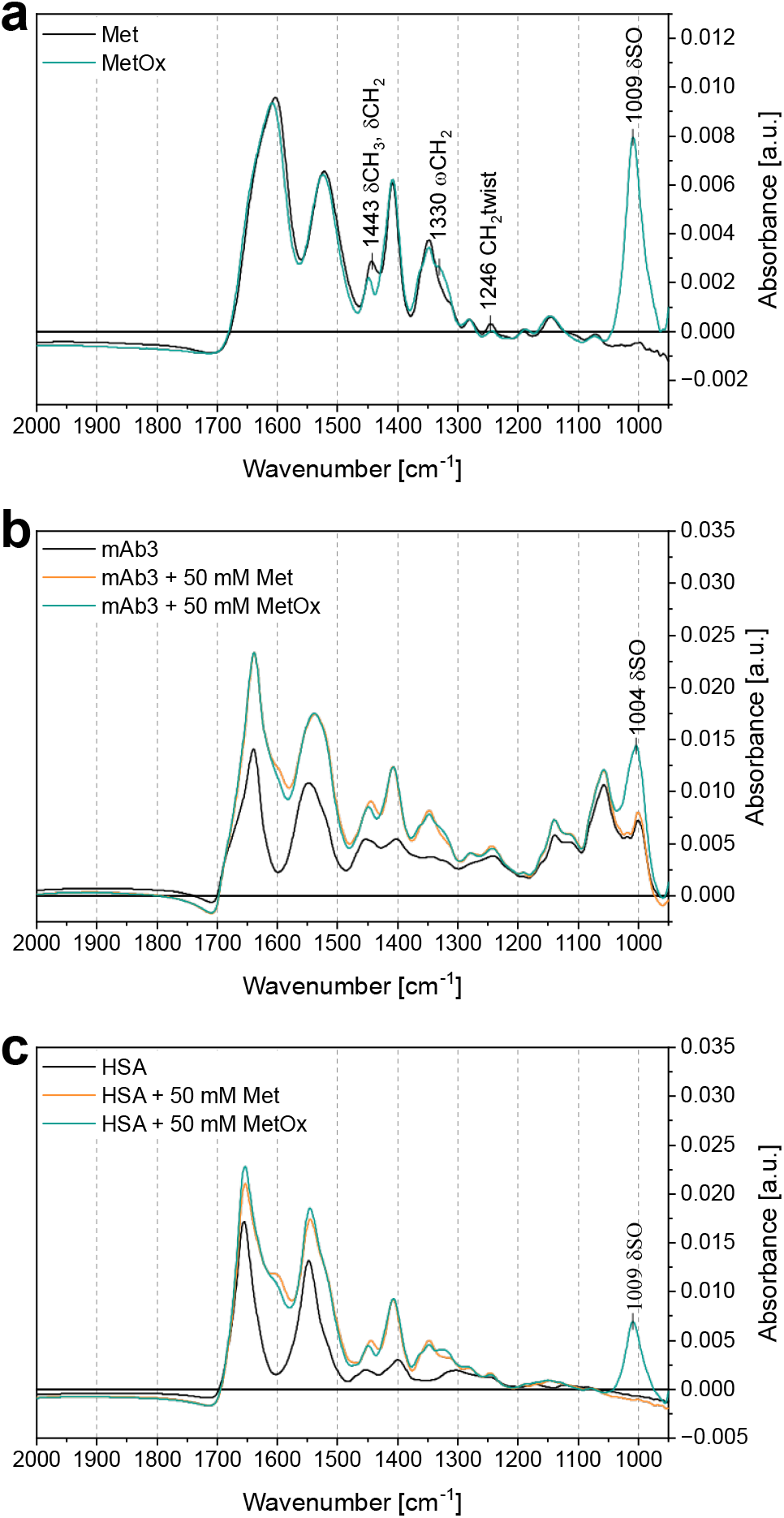
FT-IR spectra of mixtures of Met, MetOx and proteins in aqueous solution; (a) comparison of 50 mM Met and MetOx showing appearance of annotated S=O band at 1009 cm^-1^ upon oxidation of methionine; Peak assignments based on Torreggiani et al. (2011); (b) physical mixture of 10 mg/ml glycosylated mAb3 with 50 mM Met or MetOx; (c) physical mixture of 10 mg/ml human serum albumin (HSA) with 50 mM Met or MetOx.

### 3.5 Methionine oxidation in FT-IR spectroscopy – Baseline subtraction and oxidation quantification in proteins

The raw spectra need to be preprocessed to address concentration variations and instrument-related baseline shifts. The evaluation targets the S=O band at 1009 cm^-1^, originating from the oxidation of proteinogenic Met. For preprocessing, the spectrum of the unstressed protein (average of at least n = 3) is subtracted from the raw spectrum. A median concentration variation of 0.4 % (1^st^ quartile 0.2 %; 3^rd^ quartile 1 %) between replicates (98 samples with n = 3-6 replicates) and a median concentration deviation from the reference of 2.4 % (1^st^ quartile 1.0 %; 3^rd^ quartile 3.5 %) for a total of 386 single measurements was observed. In comparison, the intensity at 1009 cm^-1^ increases only by a maximum of 0.75 – 1.4 % upon oxidation of the three glycosylated proteins studied. To address concentration variations, least-variance scaling was applied to sample spectra which minimizes the standard deviation of the difference of each sample spectrum against the unstressed protein reference spectrum in the range of 1100 to 1200 cm^-1^ (140 data points). The resulting average difference in the same range was subsequently subtracted from the entire spectrum. To account for baseline curvature and improve result quality, the factor *τ* was introduced. In the employed instrument, both, water blanks and sample spectra, exhibited a random baseline drift at wavenumbers below 1000 cm^-1^. The baseline intensity *I_B_* as a function of the wavenumber frequency *f* was empirically described by the exponential term

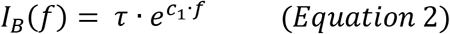

with *c*_1_ = −0.03008, determined from blank measurements. *τ* was estimated individually for each sample by least-variance regression of the baseline function (Equation 2) in the range of 950 to 960 (in selected cases up to 975) and 1100 to 1200 cm^-1^, against the difference spectrum obtained after subtraction of the unstressed reference (see also supplementary data).

### 3.6 Methionine oxidation in FT-IR spectroscopy – Linearity, Calibration and Limit of Detection

Quantification of Met oxidation via FT-IR was enabled by calibration against aqueous MetOx solutions. The peak intensity at 1007 -1009 cm^-1^ of MetOx in solution showed a high degree of linearity (*R*^2^ = 0.9995) in the tested concentration range from 0.5 to 100 mM (Figure 5 a). Consistent with the Raman assessment, the LOD of MetOx detection in FT-IR was estimated from both SNR and Equation 1. The background noise *s_B_* was approximated from the standard deviation of blank measurements in the range of 985 to 1035 cm^-1^ (n = 7). The LOD based on a 3:1 SNR was estimated at 1.63 ± 0.03 mM MetOx and at 1.97 ± 0.01 mM MetOx based on Equation 1.

**Figure 5.**
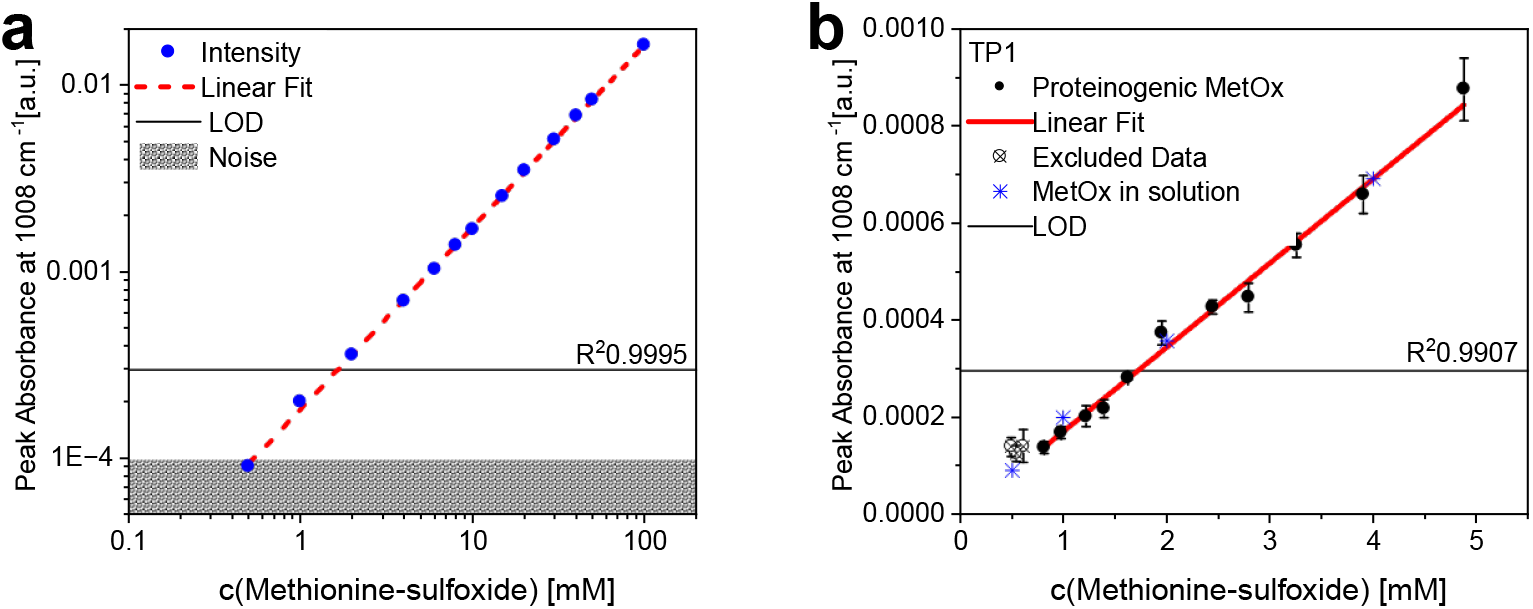
FT-IR system calibration for quantification of MetOx based on the peak intensity at 1009 cm^-1^; (a) MetOx in solution, Noise represents the standard deviation of n = 7 water baseline measurements in the range of 985 to 1035 cm^-1^, Limit of Detection (LOD) is estimated based on the 3-s_B_ criterion; (b) reference subtracted peak absorbance of a dilution series of oxidized TP1 (MetOx content calculated from MAM based on Equation 3); including linear fit and comparison to MetOx solutions.

Linearity was also observed in the detection of proteinogenic MetOx (Figure 5 b). A sample of TP1 oxidized with H_2_O_2_ at a molar ratio of 38.7 oxidant/protein, as well as unstressed reference, were diluted in matching buffer to final concentrations of 63 to 6.3 mg/ml protein. The stressed sample exhibited 42 % oxidation based on MAM measurements, corresponding to 4.9 to 0.49 mM of proteinogenic MetOx. Linearity (R^2^ = 0.9907) was observed down to 0.81 mM proteinogenic MetOx. Samples containing more than 1.22 mM MetOx showed peak absorbance significantly outside method variability. The peak absorbance of molecularly dissolved MetOx at the four lowest concentrations, as shown in Figure 5 a, correlated well with the linear fit of the proteinogenic MetOx, as shown in Figure 5 b.

### 3.7 Methionine oxidation in FT-IR spectroscopy – Proteinogenic methionine oxidation quantification and verification

Baseline- and reference-subtracted spectra of oxidatively stressed TP1 featured an increase in the peaks at 1009 cm^-1^ and at 1065 cm^-1^ (Figure 6 a). The spectral change at 1065 cm^-1^ was also observed in pure buffer under oxidative stress. Proteinogenic Met oxidation could be quantified based on the peak at 1009 cm^-1^. Importantly, the FT-IR measurement itself yields the absolute concentration of proteinogenic MetOx [mM], while MAM reports the relative percentage of oxidative changes per site. Even though both methods rely on fundamentally different measurement principles, both report the extent of methionine oxidation and can therefore be compared at the relative level of overall oxidation. The FT-IR results can be converted to the relative degree of oxidation based on Equation 3 if the total number of Met residues in the molecule *n_Met_* is known.

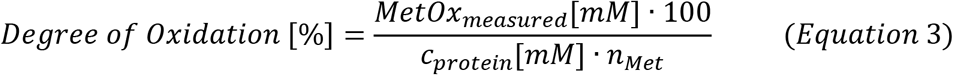

**Figure 6.**
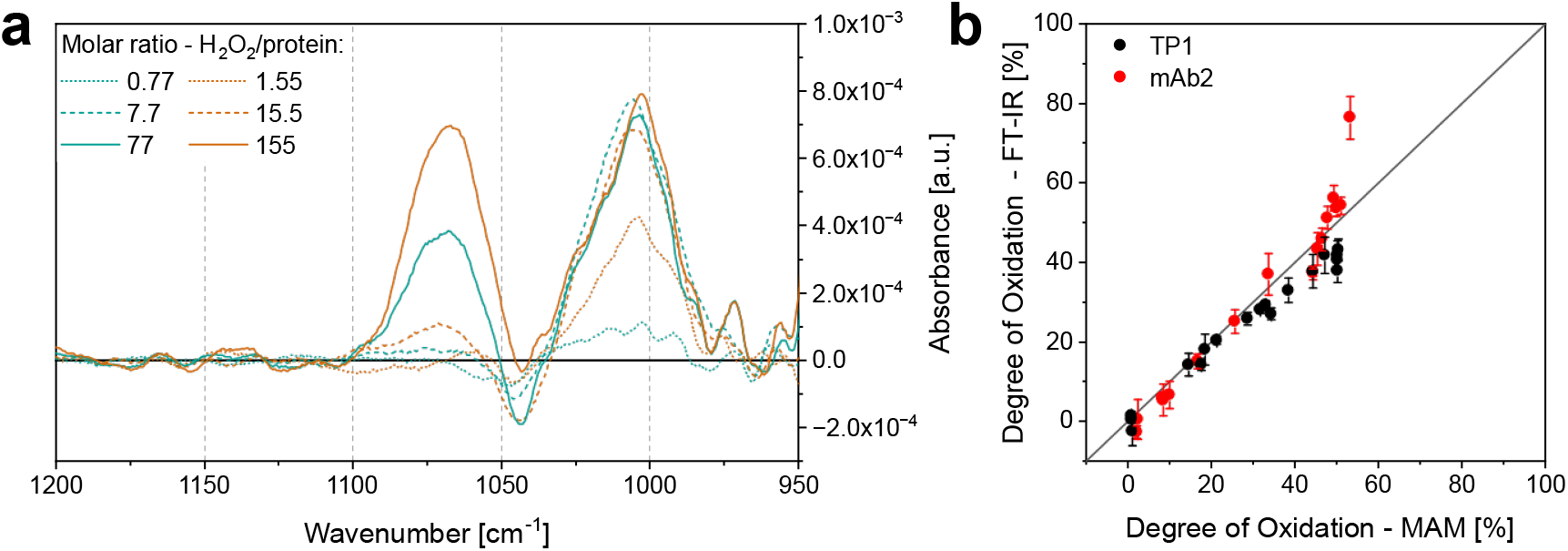
Relative quantification of Met oxidation based on the peak intensity at 1009 cm^-1^; (a) reference subtracted FT-IR spectra of TP1 at ∼ 60 mg/ml protein concentration exposed to varying amounts of hydrogen peroxide; (b) comparison of degree of Met oxidation by FT-IR and MAM for mAb2 and TP1 exposed to varying oxidative stress.

The exact *M_ww_* and *n_Met_*values of the molecules used in this study are proprietary and cannot be disclosed. Knowledge of these values does not affect the interpretation of the results or the conclusions of the study.

The degree of oxidation from FT-IR measurements was assessed in comparison to orthogonal MAM measurements and results from the two methods showed overall agreement (Figure 6 b). Up to a relative degree of ∼ 25 % of oxidation both yielded comparable oxidation levels for samples of TP1 and mAb2. For TP1 the level of oxidation at higher concentrations was underestimated in FT-IR compared to MAM. For mAb2 FT-IR and MAM results matched up to 50 % relative Met oxidation with an outlier observed at the highest degree of oxidation.

### 3.8 Methionine oxidation in FT-IR spectroscopy – Method variability

Three different preparations at four different oxidative stress levels for mAb2 and TP1 were compared (Figure 7). Series B and C were prepared in parallel on the same day, 9 months after series A. The FT-IR analysis took place immediately after incubation for all series. Series A and starting materials for series B and C were stored at -70 °C during the 9 months between series A and B/C. All samples were prepared and analyzed at the same time by MAM.

**Figure 7.**
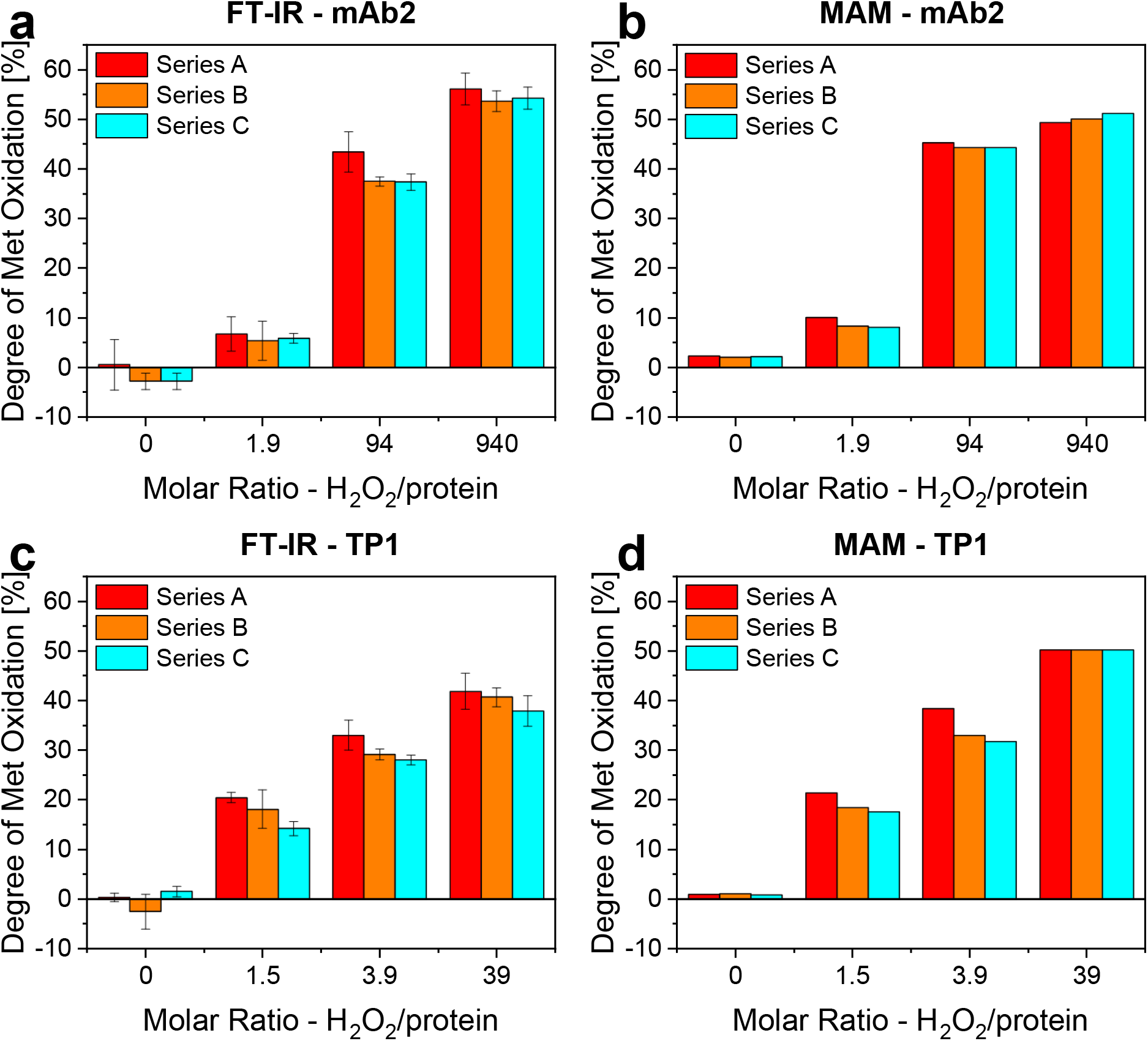
Method variability of the relative quantification of Met oxidation in FT-IR and MAM, series A was prepared 9 months prior to series B and C that were prepared on the same day, FT-IR was measured right after incubation, MAM samples were prepared together on the same day right after incubation of series B and C, series A was stored at -70 °C before analysis; (a) and (c) degree of methionine oxidation as determined by FT-IR (n = 3 measurements per sample) mAb2 (a) and TP1 (c); (b) and (d) degree of methionine oxidation as determined by MAM (n = 1 measurements per sample) of mAb2 (b) and TP1 (d).

Under all stress conditions, the variability in the measurement of the relative degree of Met oxidation was low for both molecules and with both methods. In FT-IR, the median standard deviation of the eight triplicate samples was 2.1 % with a minimum of 0.7 % and a maximum of 3.5 %. In comparison, the median standard deviation in MAM was lower, at 0.8 % (Min 0.1 %; Max 3.5 %). In several cases, series A showed a slightly elevated degree of oxidation compared to series B and C. The average variability between series observed for mAb2 was smaller than for TP1 in both methods.

## 4 Discussion

Alongside conformational, colloidal, and interfacial stability, chemical stability is a main pillar of the stability of pharmaceutical proteins. As such, it is pivotal in defining critical quality attributes and control strategies in pharmaceutical development. One of the most common chemical liabilities is the oxidation of Met residues. Peptide mapping by LC-MS-based MAM is a common analytical technique to examine the degree of oxidation and to identify oxidation liabilities of protein drug candidates on a single residue level. While it is highly sensitive, it comes with the drawbacks of extensive demand for resources and expertise. Moreover, the necessary digestive sample preparation can potentially introduce artifacts if not mitigated by preventive measures. Thus, Met oxidation detection by spectroscopic methods in native condition provides more accessible alternatives for measurements of proteins in folded state with lower resource demands in comparison to library preparation for MAM. A thorough assessment of both Raman microscopy as well as FT-IR is required to evaluate the detectability of oxidative changes in protein solutions, the feasibility of oxidation quantification and the agreement of results with orthogonal methods like MAM.

### 4.1 Suitability of Raman spectroscopy for methionine oxidation detection

Confocal Raman spectroscopy has been shown to be a powerful tool for chemical analysis and crystallinity assessment (Paudel et al., 2015). The feasibility of protein oxidation detection has been examined by Balakrishnan et al. The CS band at 702 cm^-1^ is the only spectral change specific to Met oxidation in protein samples that can be distinguished from the protein background (Balakrishnan et al., 2018). Utilizing optimized set-ups, reasonably high signal-to-noise ratios were observed for MetOx in solution. A LOD of ∼ 1.8 -3.3 mM and a linear response over the investigated range up to 100 mM demonstrated the suitability of Raman spectroscopy for quantification of MetOx in solution. The SNR of the measurements in the bulk solution had been improved by several method optimizations, including choice of laser wavelength, laser power, objective, and grating. Moreover, the use of ring slides and closed micro-chambers lowered the sample volume needed. A potential path to high-throughput, low-volume measurement of liquid solutions could be the automated scanning of a large number of closed micro-chambers in a tailor-made well-plate-style layout. Regardless of the method improvements, proteinogenic MetOx concentrations of up to 15.4 mM (mAb1; by FT-IR and MAM), well above the LOD of Raman (∼ 3.0 mM) could not be detected. The absence of the MetOx specific signal is not due to the chemical environment in a folded protein as chemical denaturation did not provide a benefit. Balakrishnan et al. have described changes related to Met oxidation in monoclonal antibodies at 50 mg/ml in a cuvette-based Raman system with a 785 nm laser (Balakrishnan et al., 2018). Regardless of the utilization of extended measurement times, elevated protein and oxidant concentrations, and optimized settings, this could not be realized in Raman microscopy, likely due to the differences in instrumentation and general measurement set-up. Thus, only the detection and quantification of MetOx in solution was possible by Raman microscopy but not of proteinogenic MetOx.

### 4.2 Suitability of FT-IR spectroscopy for methionine oxidation detection

FT-IR spectroscopy is employed in the characterization of a wide range of molecule classes. In the context of proteins, focus is usually on the amide bands for structural analysis. Previous work has shown that it can also be utilized to detect the presence of oxidative changes of molecular and proteinogenic Met (Balakrishnan et al., 2018; Ravi et al., 2011; Torreggiani et al., 2011). The most prominent change is the appearance of the S=O band at ∼ 1009 cm^-1^ upon Met oxidation. This band is well suited for oxidation detection and quantification as it does not appear in unstressed protein samples, and there are no protein-related bands that overlap with it. Their dependence on structural changes in the molecule could otherwise alter the quantification of Met oxidation based on peak intensity. Bands related to glycosylation overlay with the S=O band, but these are not sensitive to structural changes triggered by exposure to oxidative stress conditions and can be subtracted as background. Previous work employing ATR-based FT-IR set-ups for Met oxidation analysis did not investigate the feasibility of quantification (Balakrishnan et al., 2018; Ravi et al., 2011). This work demonstrates the feasibility of Met oxidation quantification in a liquid-cell FT-IR set-up, based on the distinct S=O band at 1009 cm^-1^. Linearity of the peak intensity at 1009 cm^-1^ upon dilution was observed for both MetOx in solution and proteinogenic MetOx at concentrations ranging from 100 mM down to the LOD. The estimation of the LOD was approached based on a 3:1 SNR (3-*s_B_* criterion, 1.6 mM MetOx) and on Equation 1 (1.97 mM MetOx). ICH guidelines also allow validation of lower range limits based on visual evaluation, if analytes of known, low concentration can be reliably resolved (ICH, 2023). The assessments in Figure 5 do not fulfill the requirements for this approach. Nonetheless, separation from baseline could be observed in samples of assumed proteinogenic MetOx concentration of 1.22 mM. Furthermore, several concentrations below the estimated LOD remained close to the extrapolated calibration curve. These observations justify the use of the 3-*s_B_* criterion (1.6 mM MetOx) as LOD estimate and suggest that future studies could potentially establish a lower LOD for the method. Although selected analytical performance characteristics were evaluated with reference to ICH Q2(R2), the present study cannot establish full compliance with these guidelines as it was designed as a feasibility assessment rather than a formal validation of the analytical procedure. Full validation would exceed the requirements for the intended application as an early-development screening method outside a quality-controlled environment.

The quantification of proteinogenic MetOx based on the S=O band is enabled by a newly developed background subtraction method, that accounts for baseline shifts and concentration variations. Data could be generated and evaluated quickly, accounting for about 50 samples within three days (excl. calibration), which even provides a small timely gain compared to MAM (50-60 samples in two weeks, excl. library preparation). Crucially, the relative quantification results of a monoclonal antibody and a therapeutic protein from FT-IR were in agreement with MAM results. The two techniques are in close agreement up to a level of 25-50 % oxidation. In most cases, FDS in formulation development will target lower degrees of oxidation as these are more likely to be representative of behavior upon intended long-term storage of the product. The deviations between MAM and FT-IR at higher degrees of oxidation are likely due to the increased occurrence of additional modifications not observed at lower stress level. While MAM quantifies the relative abundance of all peptides defined within its library, novel types of modifications, only occurring at extreme stress conditions beyond those included in the library generation, can cause shifts in the relative quantification. Hence, the bulk quantification in FT-IR, while being less specific, could potentially provide more accurate results at such extreme stress conditions.

Generally, the FT-IR measurements showed a low standard deviation of around 2 % for replicate measurements of the same sample. Between sample preparations, the deviations were minimal when prepared at a similar time and could only be considered relevant in one out of six cases when samples were prepared nine months apart (Figure 7). Based on this limited data set the sample preparation process, including the catalysator-mediated quenching, exhibited low variability.

The current limitation in quantifying Met oxidation in proteins by FT-IR is the need for elevated protein concentrations to ensure MetOx concentrations above the LOD. Figure 8 summarizes the concentration of proteinogenic Met in several mAbs, IgG-like bispecific antibodies (bsAb), non-canonical antibody formats (ncAb), TPs, HSA, the NIST-Fab and lysozyme at a total protein concentration of 50 to 150 mg/ml. Moreover, the amount of Met residues, that are potentially liable to oxidation was estimated based on SASA. At 50 mg/ml, all presented molecules contain Met at an abundance exceeding the LOD of the currently used instrumentation. However, for most molecules a substantial fraction of the solvent-accessible Met residues would need to be oxidized before bulk MetOx concentration exceeds the detection limit. For the MetOx concentration to surpass the LOD, even for lysozyme, ∼20 % of Met residues predicted to be liable would need to be oxidized. At higher protein concentrations, such as 150 mg/ml, this limitation plays a smaller role as the average concentration of liable Met residues is about four times higher than the LOD. Yet, proteins with naturally low abundance of Met (on the left side of Figure 8) or few oxidation-prone residues would still require a substantial degree of oxidation to enable evaluation. With a volume requirement of 50 µL per sample, an analysis at 150 mg/ml consumes a low amount of 7.5 mg protein, which is still considerably higher than the less than 0.1 mg needed for automated MAM sample preparation. In consequence, the current method is primarily applicable to FDS on high-concentration proteins containing several oxidation-prone Met residues. The sensitivity achieved with the present FT-IR setup is insufficient for routine Met stability monitoring, where even oxidative changes below 5% at individual residues can be critical. Nevertheless, the method may provide a rapid bulk-screening approach for native samples in early development, for example to define suitable oxidative stress ranges or assess formulation strategies such as antioxidant addition. Future improvements in instrument sensitivity could further expand its applicability. Given the increasing prevalence of highly concentrated therapeutic proteins, these limitations do not preclude the use of FT-IR for such early-stage screening applications.

**Figure 8.**
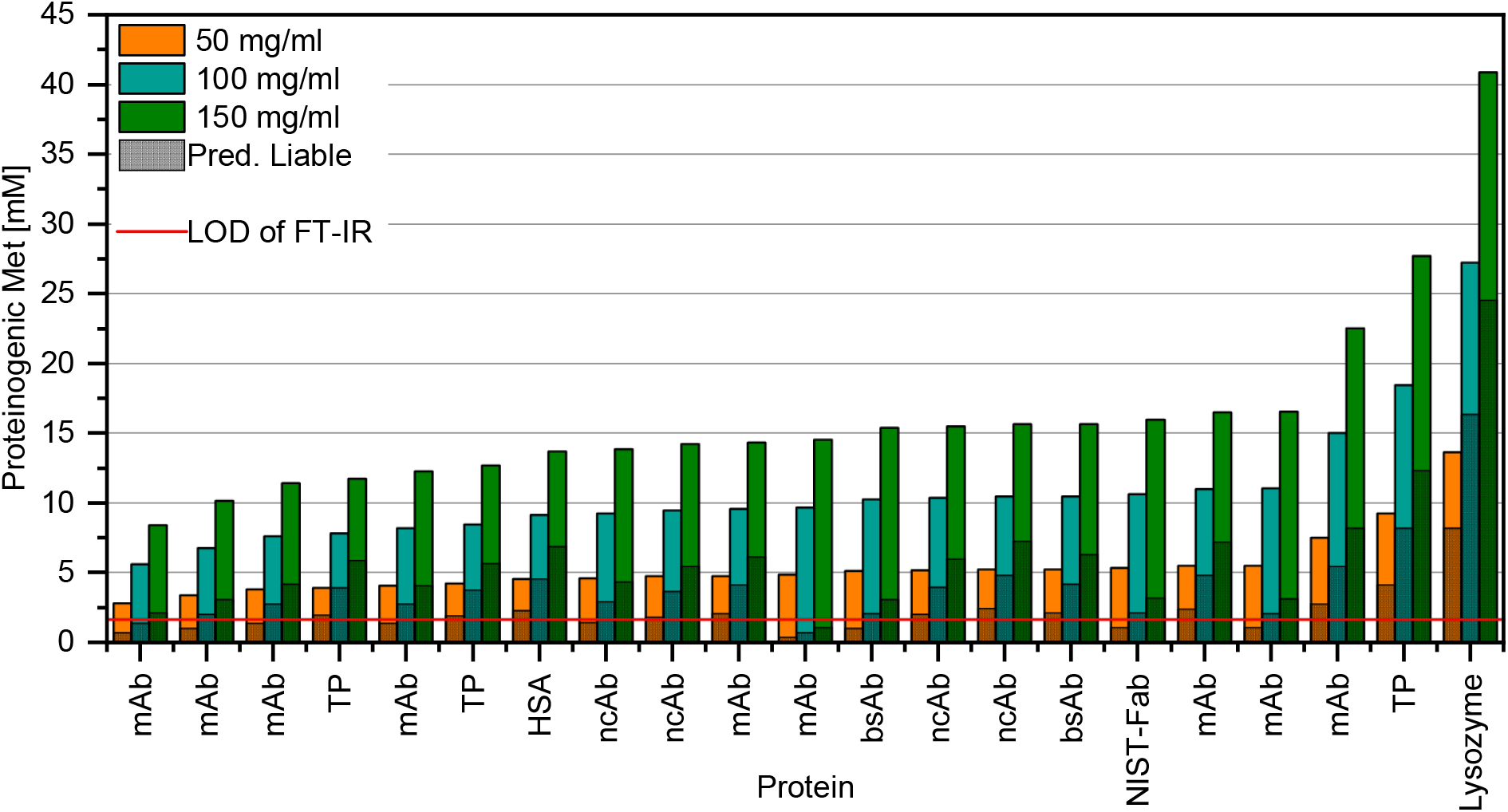
Concentration of proteinogenic methionine in mAbs, IgG-like bispecific antibodies (bsAb), non-canonical antibody formats (ncAb), TPs, HSA, the NIST-Fab and lysozyme at 50, 100 and 150 mg/ml compared to the LOD of MetOx in FT-IR as determined in this work; Shaded areas indicate Met residues that are predicted to be liable to oxidation (based on ≥ 15 % static SASA); LOD is estimated based on 3-s_B_ criterion.

## 5 Conclusion

Met oxidation is one of the most common oxidative modifications affecting the stability of protein biopharmaceuticals. Fast development timelines and the comparably higher resource demand of LC-MS methods drive the exploration of spectroscopic technologies as alternative screening tools. This work evaluated the suitability of Raman microscopy and liquid-cell FT-IR for the quantification of Met oxidation. Previous successful application of cuvette-based Raman spectroscopy proved not to be transferable to Raman microscopy. Nonetheless, the optimizations established in this study can potentially be transferred to other applications of bulk measurements in liquids.

In contrast, the findings of this work underscore FT-IR as a potential screening alternative for bulk quantification of Met oxidation in protein therapeutics. This technique does not offer the single amino acid residue resolution of MAM or the sensitivity required for routine stability monitoring in its current state and further validation would be required before application in a quality-controlled environment. Nonetheless, its ability to conduct native-state measurements and minimal sample preparation requirements position it as a valuable method, especially in early phases of development or whenever MS-based analytics are not available. The relative agreement between MAM and FT-IR results at relevant oxidation levels as determined by the data processing workflow established in this work, and the low variability increase the confidence in the potential of the method. This positions FT-IR as a promising tool for rapid, non-invasive oxidation monitoring in protein drug development.

## 6 Supporting Information

Supplementary Data: Examples of trials for detection and quantification of methionine oxidation in proteins in Raman microscopy and detailed description of pre-processing of sample spectra in FT-IR measurements.

## 7 Acknowledgments

We would like to thank Dominik Höyng from Renishaw GmbH in Pliezhausen, Germany, for the opportunity to conduct test measurements at their facilities, and for their scientific guidance on method optimization for Raman spectroscopy. Our thanks go to Nenye Johnson who supported the experimental execution of this work. Moreover, we want to express our gratitude to the Biophysical Network within drug product development at Novartis Pharma AG for their scientific guidance and Laura Truman and Thomas Pohl for their contributions in improving the manuscript.

## 8 Author contributions: CRediT

**Thorben Höltkemeier:** Conceptualization, Methodology, Validation, Formal analysis, Investigation, Data Curation, Writing – Original Draft, Writing – Review and Editing, Visualization, Project administration. **Ingo Fischer:** Conceptualization, Resources, Writing – Review & Editing, Supervision, Funding acquisition. **Marco Venerito:** Methodology, Validation, Investigation, Writing – Review & Editing. **Victor Le-Minh:** Conceptualization, Methodology, Formal analysis, Investigation, Data Curation, Writing – Review & Editing. **François Griaud:** Conceptualization, Methodology, Resources, Writing – Review & Editing, Supervision. **Patrick Merkle:** Methodology, Investigation. **Manuela Gritsch:** Conceptualization, Methodology, Validation, Resources, Writing – Review & Editing. **Adam Sobczuk:** Conceptualization, Methodology, Investigation, Writing – Review & Editing. **Malak Bitar:** Conceptualization, Methodology, Resources, Writing – Review & Editing, Supervision. **Eva Krois:** Conceptualization, Methodology, Investigation, Data Curation, Writing – Review & Editing. **Patrick Kubryk:** Conceptualization, Methodology, Investigation, Resources, Writing – Review & Editing. **Karoline Bechtold-Peters:** Conceptualization, Supervision. **Christoph Haisch:** Conceptualization, Methodology, Resources, Writing – Review & Editing, Supervision. **Wolfgang Friess:** Conceptualization, Methodology, Writing – Review & Editing, Supervision.

## Abbreviations

ATR: attenuated total reflection
bsAb: bispecific antibody
CCD: charge-coupled device
EMCCD: electro-multiplied charge-coupled device
FDS: forced degradation studies
FT-IR: Fourier transform infrared spectroscopy
HSA: human serum albumin
LC-MS: liquid chromatography coupled with mass spectrometry
LOD: limit of detection
mAb: monoclonal antibody
MAM: multi-attribute method
Met: methionine
MetOx: methionine-sulfoxide
ncAb: non-canonical antibody format
SASA: solvent accessible surface area
SEC: size exclusion chromatography
SNR: signal-to-noise ratio
TP: therapeutic protein

## 11 Tabel of Contents Graphic (TOC)

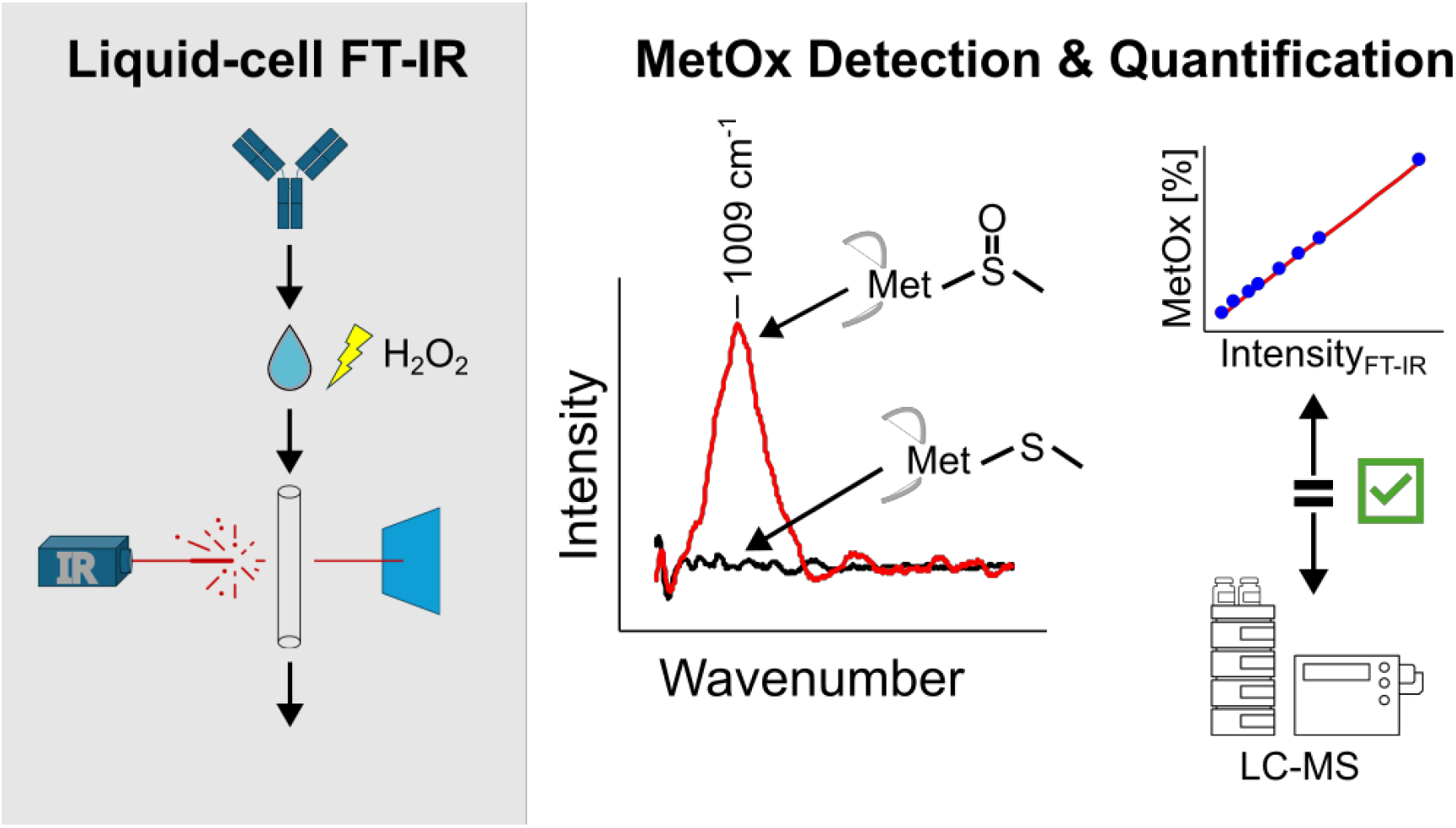

