## Supplementary Data for "Feasibility of methionine oxidation quantification in proteins through Raman and FT-IR spectroscopy"

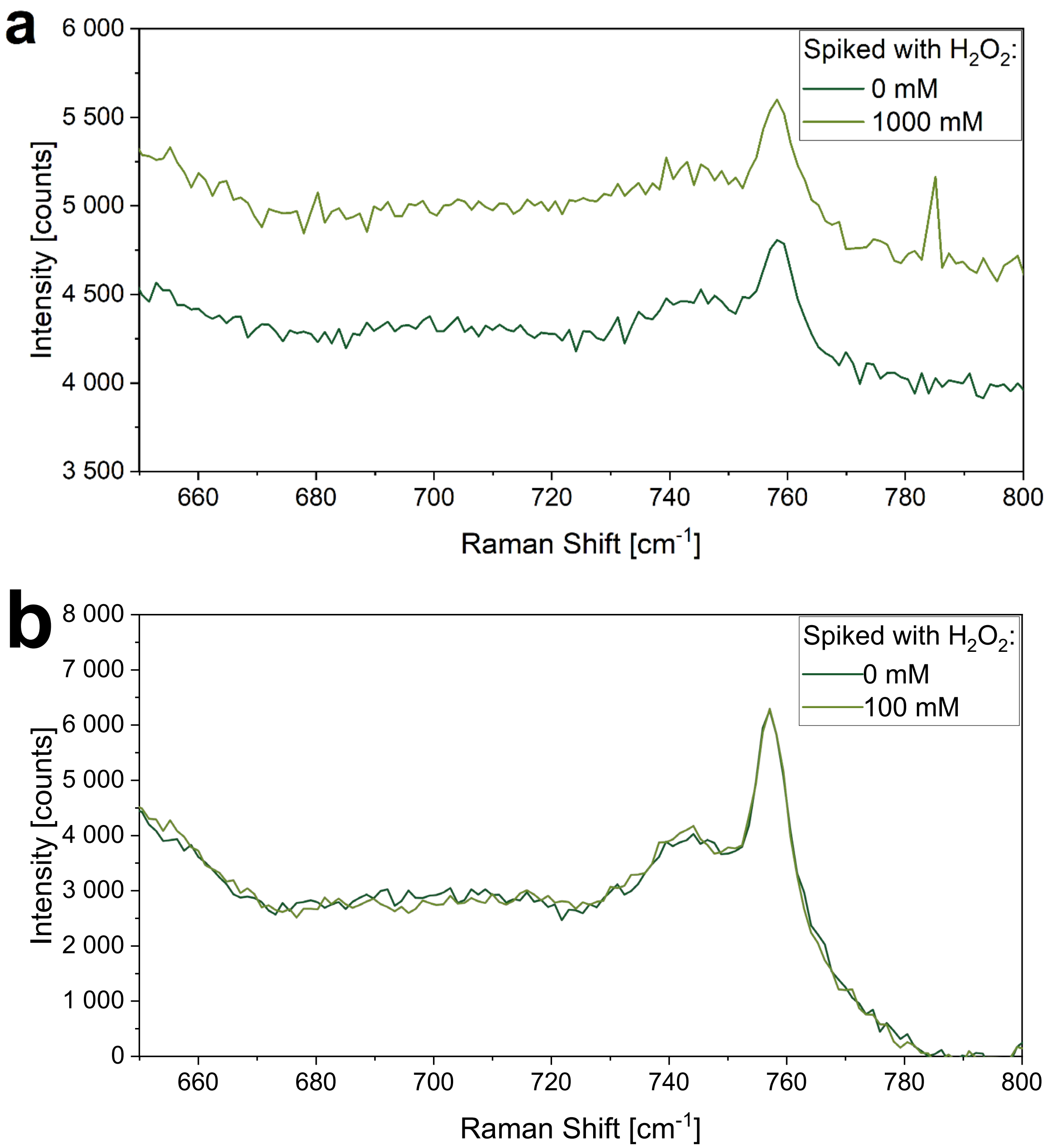

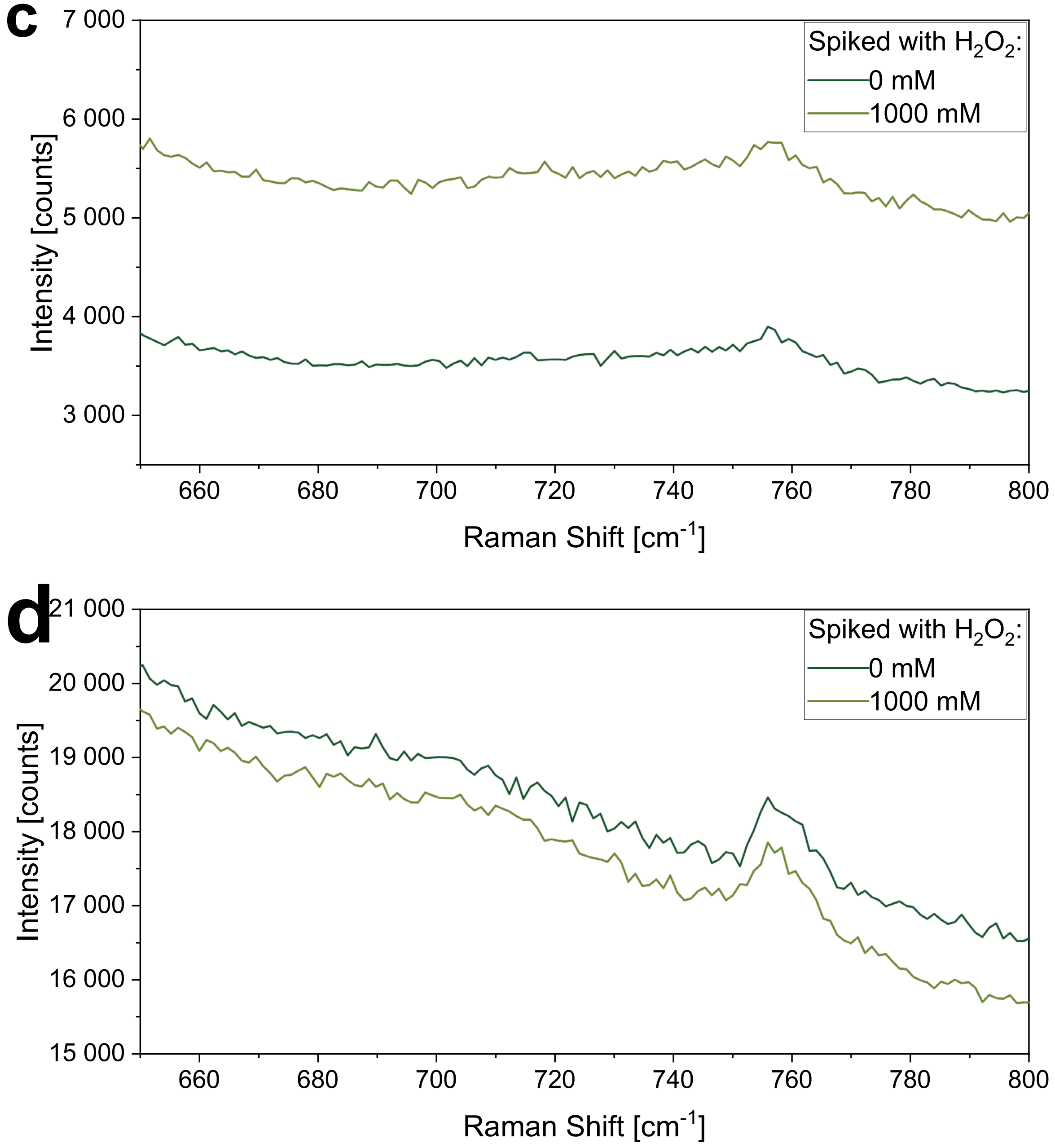


**Figure S1.** Raman spectra of Ab2 (a, 81 mg/ml), Ab3 (b, 81 mg/ml), TP1 (63 mg/ml) and TP2 (10 mg/ml) under high oxidative stress showing no change in the region of interest around 702 $cm^{-1}$. For spectra a-c the water baseline was subtracted, for spectrum d the water baseline was not subtracted.

For none of the molecules tested in this work confocal Raman microscopy did identify peaks in the area of interest around 702 $cm^{-1}$ in oxidized samples. The oxidation was proven by orthogonal FT-IR and MAM measurements. Figure S1 shows four more “worst-case scenarios" at the highest tested oxidative stress at extended measurement time of 600 s. None of the examples shows a relevant change in the area around 702 cm$^{-1}$.

*Pre-processing of sample spectra in FT-IR measurements*

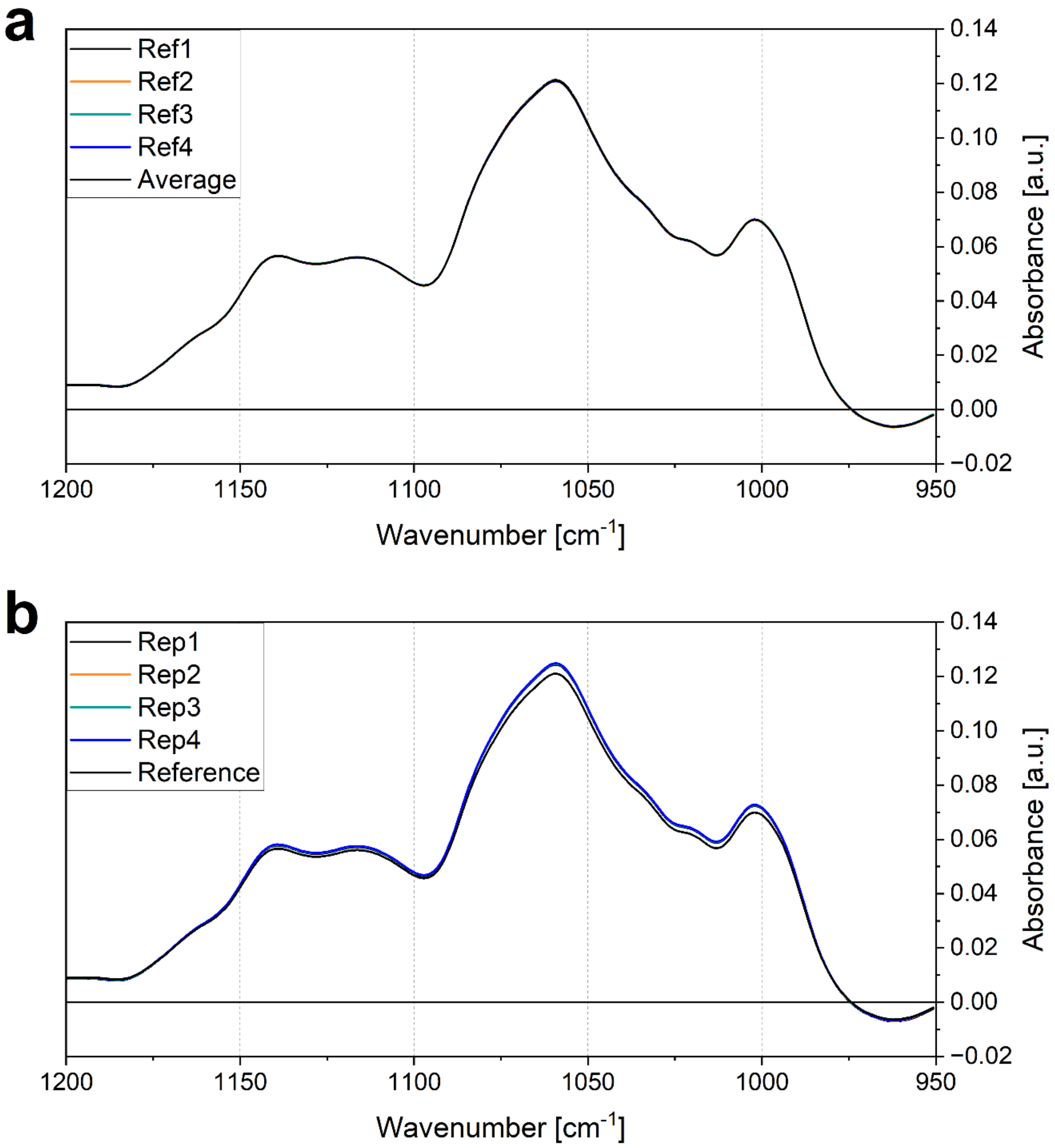

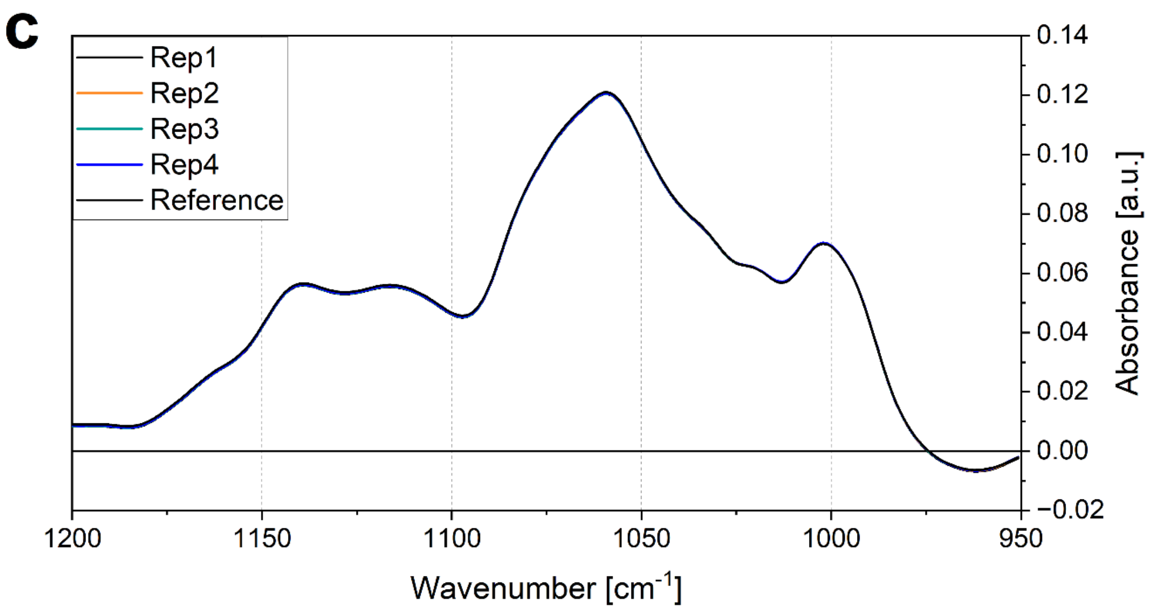


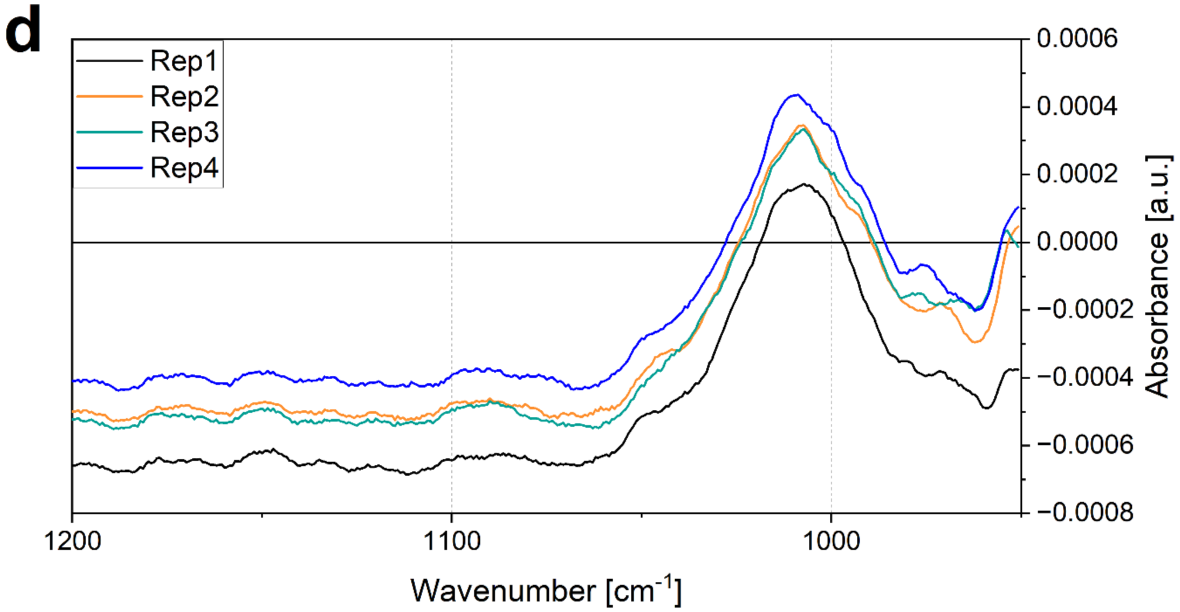


**Figure S2.** Reference subtraction in **s**pectra processing of TP1 exposed to a molar ratio of 8.7 $H_2O_2$/protein. (a) Four replicate measurements of the reference (not exposed to $H_2O_2$) and the calculated average. (b) Comparison of the reference average with four replicate measurements of the oxidized sample. (c) Comparison of the reference average with the scaled spectra of the four sample measurements. (d) Difference of the four sample spectra to the reference average after scaling.

To account for measurement variations, FT-IR spectra of oxidized samples were pre-processed.

Out of at least three reference measurements as a function of the wavenumber frequency $f$ of the unstressed sample, the average was calculated and used as reference $S_{Ref}(f)$ for further processing (Figure S2 a). Sample spectra $S_{Ox}\ (f)$ were then compared to this reference spectrum, usually showing slight variations in overall concentration due to the sample preparation process (Figure S2 b). Based on least-variance, a scaling factor α (Equation S1, Table S1) was then calculated for each spectrum individually so that the standard deviation of the difference of $S_{Ox}(f) - S_{Ref}(f)$ of all N = 140 data points in the range of 1100 to 1200 $cm^{-1}$ was minimal.

$$\alpha = \frac{\sum_{i \in F}\left(S_{Ox}(f_i) - \overline{S_{Ox,\imath \in F}}\right)\left(S_{Ref}(f_i) - \overline{S_{Ref,\imath \in F}}\right)}{\sum_{i \in F}\left(S_{Ox}(f_i) - \overline{S_{Ox,\imath \in F}}\right)^2} \qquad Equation\ (S1)$$

$$F = [1100, 1200]cm^{-1}$$

The resulting difference spectrum $S_{Diff}(f)$ is then calculated as described in equation S2 (Figure S2 d).

$$S_{Diff}(f) = \alpha \cdot S_{Ox}(f) - S_{Ref}(f) \qquad Equation\ (S2)$$

While this approach ensures a good alignment between sample and reference spectra in not being affected by baseline shifts, it results in a flat baseline with an offset as observed for the shown spectra (Figure S2 d). Consequently $S_{Diff}$ is further corrected by subtraction of the average of all data points of $S_{Diff}$ in the range of 1100 to 1200 $cm^{-1}$.

$$S_{Diff,corrected}(f) = S_{Diff}(f) - \overline{S_{D\imath ff,\imath \in F}} \qquad Equation\ (S3)$$

**Table S1** α and τ values of spectra of TP1 exposed to a molar ratio of 8.7 $H_2O_2$/protein

| Value | **Rep1** | **Rep2** | **Rep3** | **Rep4** |
|---|---|---|---|---|
| $\alpha$ | 0.9684 | 0.9656 | 0.96633 | 0.9660 |
| $\tau$ | $0.7059 \cdot 10^9$ | $1.2291 \cdot 10^9$ | $1.4611 \cdot 10^9$ | $1.2030 \cdot 10^9$ |

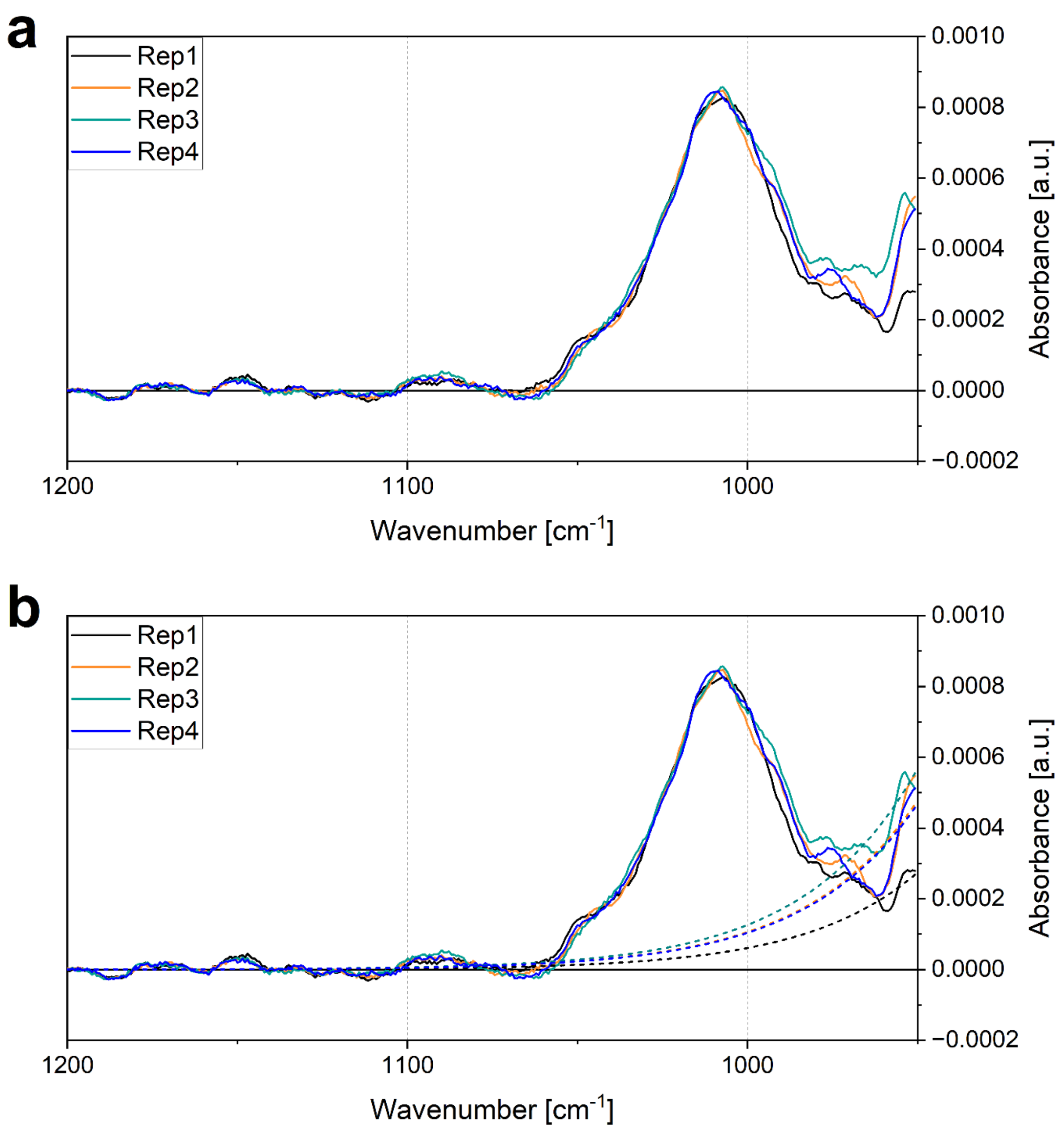
a
Rep1
Rep2
Rep3
Rep4
0.0010
0.0008
0.0006
0.0004
0.0002
0.0000
−0.0002
Absorbance [a.u.]
1200
1100
1000
Wavenumber [cm$^{-1}$]
b
Rep1
Rep2
Rep3
Rep4
0.0010
0.0008
0.0006
0.0004
0.0002
0.0000
−0.0002
Absorbance [a.u.]
1200
1100
1000
Wavenumber [cm$^{-1}$]

c

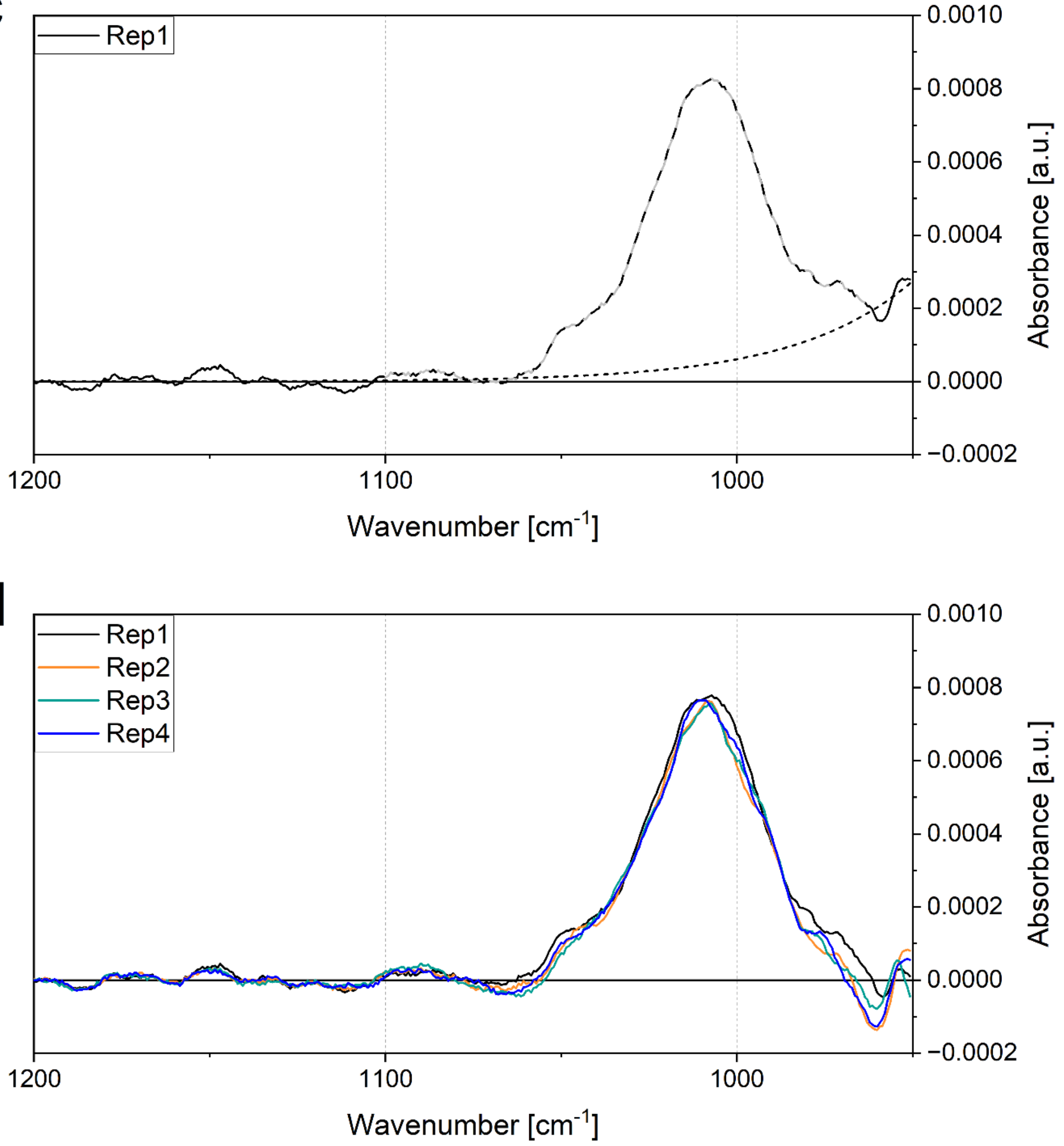


d



**Figure S3.** Baseline correction in **s**pectra processing of TP1 exposed to a molar ratio of 8.7 $H_2O_2$/protein. (a) Four replicate measurements after subtraction of the baseline offset in the range of 1100 to 1200 $cm^{-1}$ (Equation S3). (b) Same spectra as in (a) with the baseline fitted to the ranges of 950-960 and 1100-1200 $cm^{-1}$ (dotted lines). (c) Singled out representation of Rep1 from (b) with the ranges used for fitting in solid and the non-fitted range in grey-black dash. (d) Final spectra used for quantitative evaluation of proteinogenic methionine oxidation after subtraction of the baseline.

After reference subtraction and correction of the baseline offset, a distinct curving of the baseline at the lowest wavenumbers is observed that can be upward as in the example (Figure S3 a) or downward. Based on water measurements, this curving was empirically described by an exponential function with a fixed exponential factor $c_1 = -0.03008$ that describes the baseline intensity $I_B$ (Equation 1).

$$I_B(f) = \tau \cdot e^{c_1 \cdot f} \quad \textit{(Equation 1)}$$

The pre-exponential factor τ was then obtained equivalent to the scaling factor α of the reference subtraction following a least-variance approach minimizing the standard deviation of the difference between the exponential function $I_B(f)$ and the corrected spectrum $S_{Diff,corrected}(f)$ in the ranges of 950 to 960 (in other cases up to 975) and 1100 to 1200 $cm^{-1}$ (Figure S3 b & c, Equation S4).

$$\tau = \frac{\sum_{i \in G}\left(S_{Diff,corrected}(f_i) - \overline{S_{Diff,corrected,\iota \in G}}\right)\left(e^{c_1 \cdot f} - \overline{e^{c_1 \cdot f_{\iota \in G}}}\right)}{\sum_{i \in G}\left(e^{c_1 \cdot f} - \overline{e^{c_1 \cdot f_{\iota \in G}}}\right)^2} \quad Equation\ (S4)$$

$$G = [950{,}960] \cup [1100, 1200] cm^{-1}$$

The exemplary values for τ of the example spectra are summarized in Table S1.

Subtraction of the empirical exponential baseline then results in the final spectrum $S_{eval}(f)$ for evaluation (Figure S3 d, Equation S5).

$$S_{eval}(f) = S_{Diff,corrected}(f) - \tau \cdot e^{c_1 \cdot f} \quad Equation\ (S5)$$
